# An Arabidopsis root phloem pole cell atlas reveals *PINEAPPLE* genes as transitioners to autotrophy

**DOI:** 10.1101/2021.08.31.458411

**Authors:** Sofia Otero, Iris Sevilem, Pawel Roszak, Yipeng Lu, Valerio Di Vittori, Matthieu Bourdon, Lothar Kalmbach, Bernhard Blob, Jung-ok Heo, Federico Peruzzo, Thomas Laux, Alisdair R. Fernie, Hugo Tavares, Yka Helariutta

## Abstract

Single cell sequencing has recently allowed the generation of exhaustive root cell atlases. However, some cell types are elusive and remain underrepresented. Here, we use a second-generation single cell approach, where we zoom in on the root transcriptome sorting with specific markers to profile the phloem poles at an unprecedented resolution. Our data highlight the similarities among the developmental trajectories and gene regulatory networks communal to protophloem sieve element (PSE) adjacent lineages in relation to PSE enucleation, a key event in phloem biology.

As a signature for early PSE-adjacent lineages, we have identified a set of DNA-binding with one finger (DOF) transcription factors, the PINEAPPLEs (PAPL), that act downstream of *PHLOEM EARLY DOF* (*PEAR*) genes, and are important to guarantee a proper root nutrition in the transition to autotrophy.

Our data provide a holistic view of the phloem poles that act as a functional unit in root development.

## INTRODUCTION

In plants, organs originate from meristems postembrionically and are patterned by mobile signals and the positional information generated in the individual immobile cell types. Determining cell type-specific transcriptional programs is key to understanding the positional cues guiding plant development^1^. However, despite the importance of phloem in vascular plants and radial growth pre-patterning^2^, phloem gene expression is not yet well characterized. During root development, the term phloem is oftentimes used as a synonym of the protophloem sieve element (PSE), the cell type that undergoes a unique differentiation process to specialize in the transport of sap from source photosynthetic organs to distant sink tissues. This simplification is probably the result of the extensive knowledge we have about PSE specification^2, 3^ and differentiation ^4–11^. However, in the Arabidopsis primary root, the phloem pole is composed of six cells belonging to four distinct cell types: the central PSE is flanked by two phloem pole pericycle (PPP) cells to the outside and one metaphloem sieve element (MSE) cell to the inside, and both SE cells are in direct contact with the two lateral companion cells (CC) (Fig 1a).

**Figure 1.**
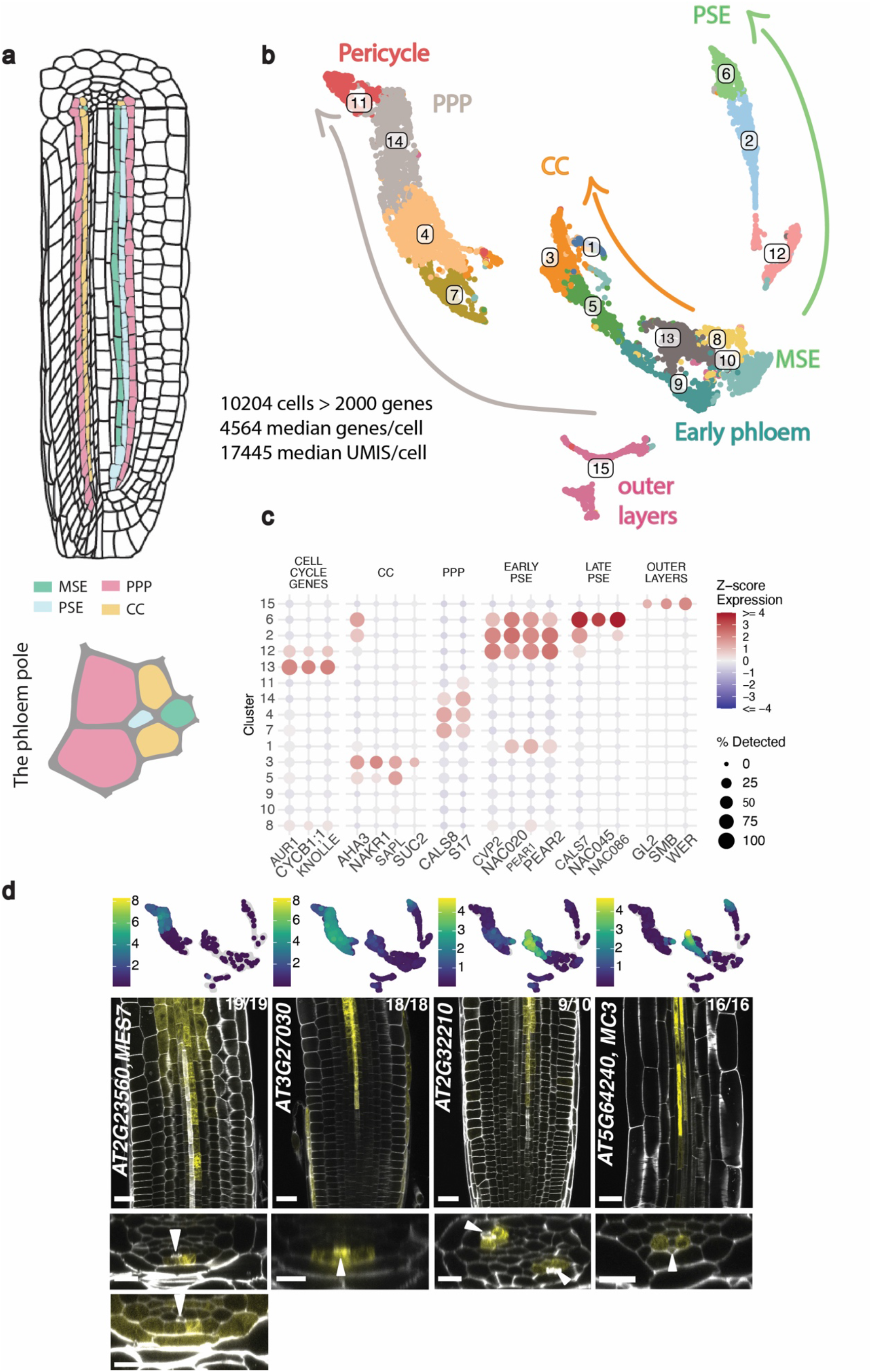
A root phloem pole cell atlas containing PSE, MSE, CC and PPP cells identified through scRNA sequencing. See next page for caption. a) Root schematic highlighting the cells in the phloem pole coloured by identity (adapted from^33^) with a close-up of the phloem pole b) UMAP plot showing the classification of 10,204 cells clustered by cell identity and developmental stage. The sample of cells has a median of 4,564 detected genes (10%- 90% percentiles: 2,600-6,780) and a median of 17,445 total UMIs per cell (10%-90% percentiles: 5941- 52689). c) Cluster annotation based on markers with known tissue- or cell-specific expression. The size of the points represents the percentage of cells in a cluster where the gene was detected (i.e. at least 1 UMI). The colour shows the scaled average expression of the gene (z-score, i.e. number of standard deviations above/below the gene’s mean across all cells). d) Newly identified genes significantly enriched in PPP (*At2g23560*, *At3g27030*) and CC (*At2g32210*, *At5g64240*). UMAPs show the particular cluster-weighted normalised expression of each gene in the phloem pole cell atlas. UMAP and microscopy pictures are representative images of the transcriptional reporter lines, where the gene promoter is fused to *VENUSer*. Scale bar in the longitudinal sections is 25 µm while it is 10 µm in the cross sections. White arrowheads point to PSE cells as a reference point. The numbers in each panel indicate samples with similar results, of the total independent biological samples observed.

In the Arabidopsis root, both conductive elements (MSE and PSE) derive from the same stem cell^12^ but MSE differentiates later, when PSE cells are no longer functional. Despite having a similar function to PSE, MSE ontogeny is less well characterized^13^ and few factors have been directly related to MSE development. An exception are the partially redundant homologs *OPS* and *OPL2*, identified as important for MSE entry into differentiation^14^. Despite some commonalities between PSE and MSE, a recent study highlighted MSE differentiation is independent of adjacent or preceding PSE^13^, underlining the peculiarities of this cell type. The conducting cell types and CC originate from different progenitors in the Arabidopsis root^12^. CC are believed to be essential to support enucleated PSE function^15^ and their intimate relationship has been evidenced by a common molecular switch controlling SE/CC fate *in vitro* and in hypocotyls^16^, while in the primary root undifferentiated CC and MSE can transdifferentiate to PSE cells if these are misspecified^17^. The CC function in leaves consists of loading nutrients into the SE but their role in the root remains elusive^18^. Traditionally, it was thought they were involved in phloem unloading^19^, that is, the exit of the nutrients from the sieve element pipe so that they reach meristematic cells for food. However, it was recently demonstrated this process happens through funnel plasmodesmata connecting PSE to PPP^20^. Despite being considered a non-vascular tissue, PPP and the associated vasculature share a high overlap in gene expression^21^ and are different in size and ultrastructure to the xylem pole pericycle (XPP) population^22^, exhibiting specific gene expression^23^ from early stages, mirroring the diarch pattern in the Arabidopsis vasculature^24^.

In the last 15 years, transcriptomics has been the stepping stone to learn about plant organogenesis. However, even if markers for mature CC and PPP were used for transcriptomics^1, 25, 26^, the lack of specific markers for early phloem, combined with the difficulties to access phloem cells, deeply embedded in the root cylinder, have hampered the study of these populations, oftentimes masked under the concept “stele”, that groups pericycle and vasculature ^27–29^. The more recent root single-cell atlases confer a detailed root panoramic but even here phloem cells remain underrepresented compared to more accessible root layers^30–32^.

Combining fluorescent activated cell sorting (FACS) and SMART-seq single cell technologies allowed the profiling of PSE at an unprecedented resolution^33^. However, PEAR genes, architects of early phloem development, do not act exclusively in PSE: they move from PSE to neighbouring cell types where they target multiple genes^2^, activating periclinal cell divisions and other unknown processes. Therefore, to gain a better understanding of phloem we need to profile all the cell types in the tissue in relation to each other along the root. We have generated a phloem pole cell atlas by sorting phloem marker lines combined with single cell sequencing, allowing us to gain resolution in cells that are underrepresented in general root cell atlases. We investigated not only the specificities of each cell type but also the transcriptional commonalities between them. We additionally identified a second set of DOF transcription factors (TF) expressed in the PSE adjacent cells, target of PEAR TF, that are important in the transition to autotrophy in young seedlings, linking phloem development and root physiology.

## RESULTS

### Cell atlas of the phloem pole

In order to profile phloem cells, we took advantage of new and existing fluorescent markers expressed in SEs, CC and PPP from early meristematic cells until differentiation (Fig S1a). This allowed us to enrich our data with cells of interest, by using FACS and preparing single-cell sequencing libraries using the 10x Chromium droplet-based protocol. This resulted in a total of 10,204 high-quality cells, defined as those having at least 2000 detected genes and no more than 10% of reads assigned to mitochondrial genes (the resultant sample of cells had a median of 17,455 reads/cell and a median of 4,564 genes/cell). The raw count data was normalised using variance stabilising transformation^34^ and integrated across batches using the mutual nearest neighbours algorithm^35^, although our main conclusions are robust to normalisation and batch effects. These cells were grouped into 15 clusters using the Louvain algorithm on a shared-nearest-neighbour cell graph and visualised using uniform manifold approximation and projection (UMAP)^36^ (Fig 1b). Using signature marker genes (Fig 1c) we identified all the cell types included in the phloem pole. One main branch corresponds to PSE conducting cells (clusters 12, 2, 6), another to CC (clusters 5, 3, 1) and a third to PPP (clusters 7, 4, 14, 11), all emerging from a central group of meristematic cells (clusters 8, 9, 10, 13). Clusters 10 and 1 express MSE genes. In turn, clusters 13 and 12 contain G2/M cell cycle markers, indicating cells undergoing division. While it is hard to distinguish any identity in cycling cells, cluster 12 expresses PSE markers, therefore probably pointing towards PSE dividing cells. Finally, cluster 15 corresponds to the outer layers of the root, as an apparent contamination during cell sorting.

Separated from the rest, clusters 7, 4 and 14 were contributed to mainly by *pS17::GFP* and *pAPL::3xYFP* markers (Fig S3a), and expressed genes characteristic of PPP such as *S17* (*At2g22850)* and *GLUCAN SYNTHASE-LIKE 4* (*CALS8, At3g14570)* (Fig 1c). In turn, cluster 11, mainly contributed to by *pS17::GFP* and the *pMAKR5::MAKR5-3xYFP* sortings, represents mature pericycle cells, since in addition to PPP markers it also expresses markers for XPP (*At1g02460*, *At4g30450*^37^, *At2g36120*, Fig S3c) and PPP (Fig 1c). This is likely because *MEMBRANE-ASSOCIATED KINASE REGULATOR 5* (*MAKR5, At5g52870)* is expressed in the whole pericycle layer high up in the root and pericycle cells come together with PPP cells for similarity.

Considering genes that were statistically more highly expressed in PPP-specific clusters, we built reporter lines for two genes, which were confirmed to have PPP-specific expression. One of these, *At3g27030*, was expressed in PPP and late PSE, while the other, *METHYL ESTERASE 7* (*MES7, At2g23560),* was expressed early in PPP and soon afterwards becomes more broadly expressed in the vasculature and endodermis (Fig 1d).

In turn, the known CC genes are expressed in cluster 5 (*SISTER OF APL*, (*SAPL, At3g12730*^20^)), with clusters 3 and 1 expressing mature CC genes (*ATPase3* (*AHA3, At5g57350*^2^), *SODIUM POTASSIUM ROOT DEFECTIVE 1* (*NAKR1)*^38^, *SUCROSE PROTON SYMPORTER 2* (*SUC2, At1g22710*^39^). *AHA3* in particular was statistically more highly expressed in this cluster and allowed the discovery of new CC genes by correlation, which were validated building reporter lines (Fig 1d). One of these was *At2g32210*, which is expressed first in PSE and then switches to a strong CC-MSE expression. In turn, *METACASPASE 3 (MC3, At5g64240),* was expressed in late PSE and started being expressed in CC after enucleation, first in a patchy way and then getting continuous and CC- exclusive. Cloning reporter lines for other genes expressed in these clusters, we found a gene expressed in PSE and CC (*PHOSPHATIDYLINOSITOL-SPECIWC PHOSPHOLIPASE C5 (PLC5, At5g58690*)), previously described to be expressed in vascular tissues^37^ and *At2g38640*^40^ mostly specific of mature CC (Fig S1b). Therefore, we have been able to validate our UMAP finding genes more enriched in the different cell types.

### Spatiotemporal patterns of phloem differentiation in the atlas

From our initial cell annotation, it seemed clear that our data also captured the temporal aspect of cell differentiation in the phloem. For example, marker genes usually expressed in more differentiated cells, showed higher expression at the tip of the “branches” on our UMAP projection, while those closer to the cycling cells could be meristematic. To validate this hypothesis, we compared our data with root longitudinal sections from previous bulk transcriptomics^1^, assigning each of our cells to the longitudinal section with which they had the highest Spearman correlation (Fig S2a). Using this strategy, we observed that the cells towards the centre of our UMAP matched with the meristematic sections of Brady *et al.*, with a temporal progression in every “branch” of our UMAP, until the more mature cells cap each trajectory. This analysis validates our hypothesis of a temporal trajectory that is well captured by our UMAP projection and cell clustering.

To further infer developmental trajectories and order our cells along a continuous pseudotime, we used Slingshot^41^ (Fig 2a). Setting a unique origin for all in cluster 13 (cycling cells), we obtained 5 different trajectories (Fig 2a), reflecting the known developmental trajectories in the root. Furthermore, these trajectories agreed with RNA velocity analysis using scVelo^42^, with velocity vectors aligning towards the end of these trajectories (Fig S2c).

**Fig 2.**
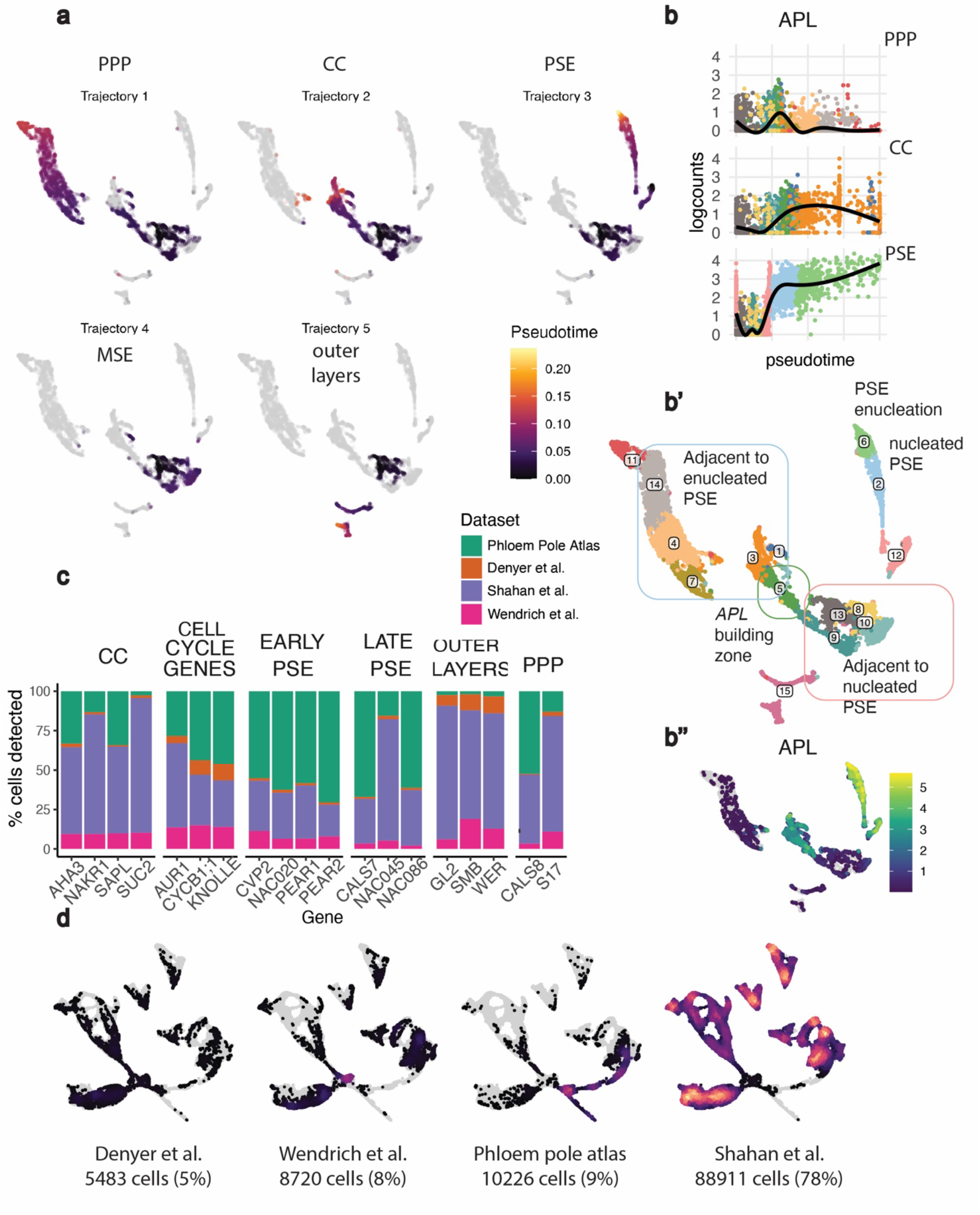
A continuous phloem developmental gradient allows the inference of developmental trajectories, which facilitate mapping of the PSE enucleation point. See next page for caption. a) Developmental trajectories inferred using Slingshot coloured according to pseudotime, with more mature cells in yellow. The origin for all trajectories was set in the clusters containing cycling cells. b) *APL* expression is plotted along the PPP, CC and PSE trajectories, with the cells coloured by cluster number in the UMAP. b’) *APL* is used as a standard to coordinate the three trajectories. Cluster 5 groups the cells with an increasing expression of *APL* in PPP and CC trajectory, mapping the enucleation point in the adjacent cells. The position of each cell type is indicated in the UMAP in relation to PSE enucleation b’’) UMAP of APL expression in the phloem pole cell atlas c) Bar plot representing the percentage of cells expressing the markers listed in the x-axis of the integrated UMAP containing 113,240 cells. Colours indicate the single cell project. d) Integrated UMAP with the cells contributed by each individual project plotted on top (number indicated below, percentage of the total in brackets), using a coloured scale to indicate cell density.

Trajectories 1-3 account for PPP, CC and PSE respectively. Trajectory 5 is for outer layers and we will not focus on it.

While PSE trajectory is independent from all others, PPP and CC have cluster 5 in common. While other clusters were unequivocally assigned to a single trajectory (see for instance cluster 3, with a probability of 1 to belong to CC trajectory, or cluster 4, with a probability of 1 to belong to PPP trajectory, Fig S2e) or shared by all trajectories (like early phloem cells in cluster 8, Fig S2e), cluster 5 was not a clear cut, with 75% chances of belonging to trajectory 2 (CC) and 25% chances of belonging to trajectory 1 (PPP), Fig S2e.

Regarding gene expression, cluster 5 has 64% cells expressing the CC marker *SAPL* (409/638 cells) and 20% expressing the PPP marker *S17* (127/638 cells), with 12% of the cells in this cluster expressing both genes simultaneously (76/638 cells). This indicates more cells in cluster 5 express CC markers than PPP markers. This matches our observations in the root, when *SAPL* starts to be expressed earlier in development than S17 (Fig S2g). The fact that a small percentage of cells express both markers at the same time despite being specific for different cell types would indicate transcriptional reporters are not always highlighting weak gene expression, so it is possible that our transcriptional data paint broader expression domains than the ones visible with the specific marker lines (see for example the broader *SAPL* expression domain compared to the cells sorted using p*SAPL::VENUSer* reporter line, Fig S3a,b).

There is no PPP-specific known marker gene expressed earlier in root development that would facilitate separation of CC and PPP identities. However, there should be an intermediate developmental stage in PPP provided by “MAKR5 sorting including one third of the root”, which we interpret as cluster 5. These cells would sit in between the early PPP cells, sorted using *MAKR5* reporter enriched in root tips, and those expressing *S17*, sorted using *pAPL::3xYFP* and *pS17::GFP* markers.

Therefore, despite different origins, a cluster common to CC and PPP trajectories points to a common transcriptional cell state between the precursor cells of these two cell types before they differentiate.

The developmental trajectories obtained reinforce clusters 8, 9 and 10 as early CC, PPP and SE cells. Given these populations are contributed mainly by cells sorted using *MAKR5* and *PEAR1del* (Fig S3a), we can conclude that these clusters correspond to the early phloem cells, containing three different identities (MSE, PPP and CC), still undifferentiated.

There is no gene statistically enriched in cluster 9 and those few in cluster 8 (Supp Table 1) are broadly expressed in whole root single cell data. Except for PSE, when we detect cycling cells expressing PSE markers in cluster 12, it is hard to distinguish an early identity in the other trajectories. However, when early phloem cells are compared to the early cells in general root cell atlases, phloem early cells cluster together more than expected by chance compared to other early cells, suggesting early phloem cells have a specific signature (Fig S4c,d).

An important event in phloem development is the enucleation of PSE, since at that moment this cell type loses the nucleus and stops directing phloem progression, becoming dependent on neighbouring cells for survival and probably triggering changes in their transcriptomes. In order to map the enucleation point in the UMAP and know which cells are neighbouring PSE before and after enucleation, we needed to coordinate trajectories, since each trajectory has a different pseudotime. To coordinate them we used our knowledge of *ALTERED PHLOEM DEVELOPMENT* (*APL*) expression, which is expressed at different times in all three trajectories, combined with the enucleation markers *NAC DOMAIN CONTAINING PROTEIN 86* (*NAC086, At5g17260)* and *NAC45/86-DEPENDENT EXONUCLEASE-DOMAIN PROTEIN 4* (*NEN4, At4g39810)* (Fig 2b, Fig S2f). *APL* is first expressed in PSE and at the time of enucleation is transcriptionally activated in CC and MSE^4^. In reporter lines like *pAPL::3xYFP* we perceive a strong signal in PPP as well (Fig S1a), a circumstance that we took advantage of for sorting, but the phloem pole cell atlas transcriptomics data do not reflect *APL* expression in mature PPP (Fig 2b”, Fig S2d). In another reporter line, *pAPL::YFPer* we also observe a signal in PPP that gets weaker going shootward (Fig S2d). Based on reporters and transcriptomics data, the signal in PPP is probably not the product of gene expression in this cell type but likely caused by direct unloading from PSE that lags and gets diluted in successive cell divisions.

While *APL* expression increases in PSE trajectory until enucleation, it starts being detected in PPP and CC trajectories in the common cluster 5 (Fig 2b), indicating this is the transition zone when *APL* starts building up in the neighbouring cell types before enucleation. Therefore, the PSE trajectory is contemporary to the early phloem cells, cluster 5 coincides with PSE enucleation preparation and clusters for mature PPP and CC contain cells that are neighbouring an enucleated PSE (Fig 2b’).

### Trajectory and marker gene analysis help identify first stages of MSE development

MSE is difficult to identify since there are no specific markers available for this cell type. However, sorting using marker lines reduces the possibilities for the output, conferring us an advantage to identify the most unknown populations over the whole root cell atlases, when possibilities for unknown clusters multiply. Slingshot identified an incomplete trajectory (trajectory 4, Fig 2a) that is mainly formed by cluster 10 (Fig S2b), which is mostly contributed by *MAKR5* sortings, pointing this could be early MSE cells (Fig S3a). In cluster 10, we find cells expressing MSE markers like *sAPL*, *APL* (Fig S3b), and other genes expressed in MSE and other cell types but excluded from PSE (*At5g47920*, *PAPL1*, Fig S3b, Fig 3b, Fig 4).

**Fig 3.**
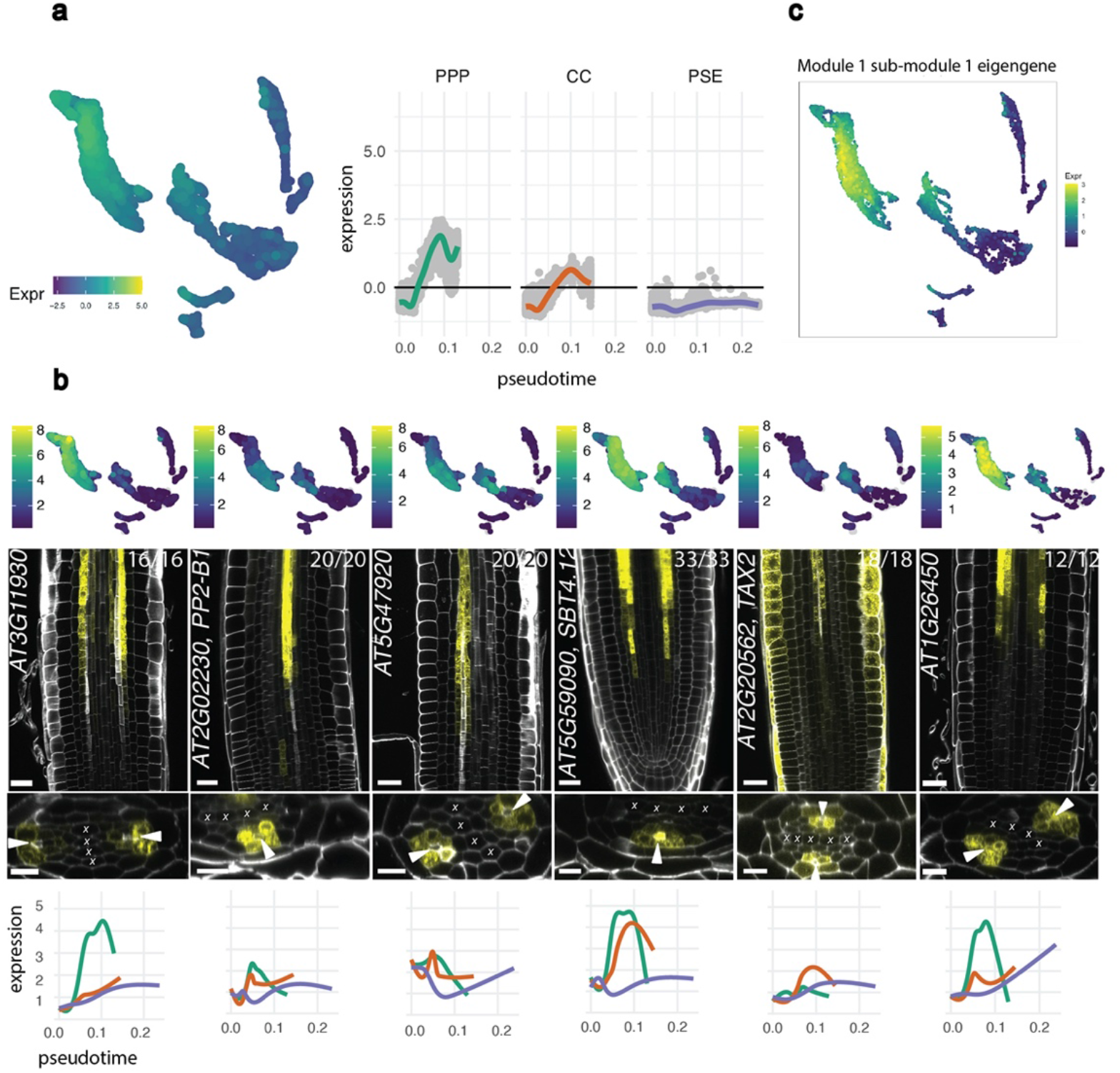
Gene regulatory network analysis identifies a gene expression pattern common to non- PSE cells that is frequent after PSE enucleation. a) The module 1 eigengene profile with its expression along PPP, CC and PSE trajectories. b) New genes with an expression pattern validating the gene profiles grouped in module 1. All the genes presented in this panel are expressed forming a ring around PSE at the time of PSE enucleation. *SBT4.12* and *TAX2* are also expressed in late PSE, with *TAX2* also showing expression in the epidermis. UMAPs show the particular cluster-weighted normalised expression of each gene in the phloem pole cell atlas and microscopy pictures are representative images of the transcriptional reporter lines where the gene promoter is fused to *VENUSer*. Scale bar in the longitudinal sections is 25 µm while it is 10 µm in the cross sections. White arrowheads point to PSE cells as a reference point. “X” marks xylem cells. Each gene has also been plotted in PPP (green), CC (orange) and PSE (purple) Slingshot trajectories, showing average expression values in the Y-axis and pseudotime in the X-axis. c) Expression profile of sub-module 1 eigengene, the sub-module of module 1 which is enriched for genes with ring-specific expression. This sub-module contains all the genes in the panel except for *TAX2*, which was not present in our network. The numbers in each panel indicate samples with similar results, of the total independent biological samples observed.

**Fig 4.**
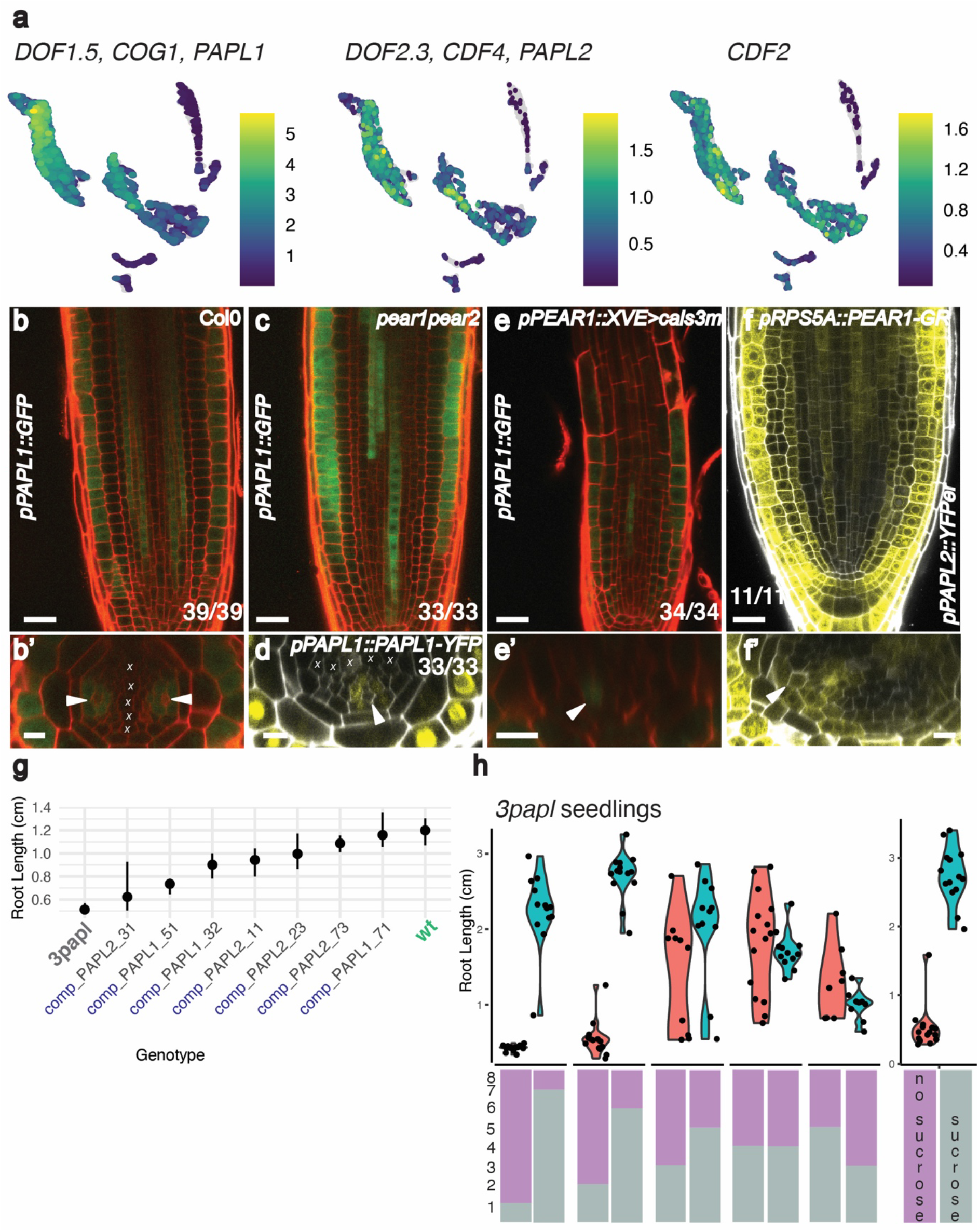
*PAPL* genes are *PEAR* targets forming a ring since the early phloem and influence root nutritional status. a) UMAPs showing the cluster-weighted normalised expression of *PAPL1*, *PAPL2* and *CDF2* in the phloem pole cell atlas. Note these genes are also expressed in the early phloem cell clusters. b) *pPAPL1::GFP/GUS* expression domain. This gene is expressed in all the cells around PSE from 40 µm from the QC and in the epidermis c) The phloem expression of *pPAPL1::GFP/GUS* is delayed until 120 µm in *pear1pear2* mutant background d) *pPAPL1::PAPL1-YFP* (Col0) translational domain coincides with the transcriptional domain e) The ring pattern of *pPAPL1::GFP/GUS* gets distorted upon PSE plasmodesmata closure using the *cals3m* tool (*pPEAR1::XVE>>cals3m*) f) *PAPL2* (*pPAPL2::VENUSer*) becomes ectopically expressed upon *PEAR1* overexpression in the meristem (*pRPS5A::PEAR1-GR*) g) Overall average root length of 6 days post-sowing (dps) seedlings of *3papl*, wt and complementation lines in *3papl* background with genomic constructs for *PAPL1* (3 lines) or *PAPL2* (4 lines) in sucrose- depleted media across experiments. The points denote the median and error bars the 95% confidence interval estimated by bootstrap (500 samples), accounting for experimental and seed stock batches. The non-parametric bootstrap approach was used due to the bimodal distributions observed in some experiments. N = 24 - 46 (median 36) seedlings per experiment. h) Transfer experiment between sucrose and sucrose-depleted plates of *3papl* seedlings. Time (in days) spent in sucrose and without sucrose is represented by a grey and purple bar respectively. Transfer was done days 1-5 and all roots were measured at 8 dps. The bars are divided in 8 portions representing the days in each condition. 539 seedlings were measured, 131 for *3papl* control (original stock), 135 for wt control (stock1) and an average of 24 (*3papl*) and 29 (wt) total seedlings per transfer experiment In confocal pictures b, e and f, primed letters show the cross sections of each respective letter. Scale bar in the longitudinal sections is 25 µm while it is 10 µm in the cross sections. White arrowheads point to PSE cells as a reference point and “X” marks xylem cells. The number in each confocal picture indicates samples with similar results of the total independent biological samples analysed.

In addition, we know procambial markers like *PIN-FORMED 4* (*PIN4, At2g01420)* become excluded from MSE cells early in development^33^ and we find this marker absent from cluster 10 (Fig S3b). Cluster 10 is mostly statistically enriched in S-phase genes (histones), indicating these are still early cells and therefore harder to characterise further.

We also know that MSE cells should not display PSE markers in early stages, since these are no longer expressed in MSE after lineage bifurcation, but will express these signature genes later in development. For this reason, we interpret that cluster 1 is a more developed MSE, since we find early SE genes like *PEAR1* and *S32* (*At2g18380*) expressed in this cluster, which is mainly contributed by the APL sorting. Cluster 1 belongs to CC trajectory, possibly because CC and MSE at this stage share some transcriptional expression, as evidenced by reporters like *At2g32210* (Fig S3b), *SAPL* and the cases shown in Fig 3b, highlighting how phloem pole cell fates are intertwined along development.

While we have end points for our PSE, CC and PPP trajectories, we don’t expect to have an endpoint for MSE, since this cell type differentiates further away from the meristem^13^. Out of the 7 genes identified to be expressed in both sieve elements^13^, we detected all the genes in PSE clusters but only *DESIGUAL2* (*DEAL2, At4g21310)* in cluster 1 and CC, confirming we have not sampled mature MSE cells. However, we are convinced we have identified the first stages of MSE development in clusters 10 and 1.

### Root phloem pole cell atlas represents a continuum of phloem development relevant for the whole plant

In order to explore the depth of our data, we integrated our phloem atlas with existing root single cell datasets^30, 32, 43^ (Fig S4a). After filtering, this process rendered a UMAP with 113,340 reclustered cells, of which 9% belonged to our project, 7.69% to Wendrich et al., 4.84% to Denyer et al. and 78.4% to Shahan et al. Despite our modest contribution regarding total cell number, we are contributing more than 50% of total cells expressing PSE and PPP markers (CALS8), and more than 25% of cells expressing CC markers (Fig 2c).

In addition, we are adding approximately one third of cycling cells (Fig 2c), indicating we have extensively profiled early cells. Interestingly, when looking at the UMAP, a continuity can be observed in the cells contributed by our atlas covering the gaps in the other data (Fig 2d). Indeed, most of the intermediate PPP and CC cells were provided by our dataset, demonstrating the difficulty to sample this group without using an active strategy to enrich this population. The continuity observed in the UMAP allowed us to track phloem developmental trajectories more accurately (Fig 2d) and enrich populations that were underrepresented in other general root atlases.

We also wanted to compare root phloem with a recently published single cell dataset on leaf, containing 478 vascular cells^44^. In Arabidopsis leaves (Fig S5a), veins are often formed by multiple sieve elements usually surrounded by at least two CC and one phloem parenchyma cell. In turn, phloem parenchyma cells, which are more irregular and have a much less dense cytoplasm compared to CC, are often in contact with one or more CC, sharing comparatively many more connections than other interfaces^45^.

When the root and leaf data were integrated, PPP and phloem parenchyma cells blended in two clusters (Fig S5b,c). Cluster 9 of the integrated data was formed by CC cells, which are present in both leaf veins and roots. However, cluster 6 of the integrated data contained a mixture of cells annotated as mature root pericycle cells and phloem parenchyma cells from leaves, showing expression of signature PPP (Fig S5e), XPP (Fig S5g) and phloem parenchyma genes (Fig S5f) in both datasets. Phloem parenchyma leaf genes are expressed in clusters 11 and 14 in the phloem pole cell atlas, corresponding to pericycle and PPP respectively (Fig S5f). Pericycle tissue is present in roots and stems, but not in leaves, and phloem parenchyma cells are found in the aerial tissue and root secondary phloem but they are not found in the primary root. As in the root, vein anatomy and cell position seem to be crucial for cell identity. Despite being different cell types with different origins, the transcriptional overlap between phloem parenchyma and mature pericycle is another indication of the importance of positional information for cell function in plants, reinforcing the role of PSE as phloem organiser. These data also suggest parenchymatous cells share similarities across different organs and underscore their relevance for phloem.

### Cells in the phloem pole share transcriptional programmes despite individual differences

In order to identify groups of genes showing distinct expression patterns in the phloem poles, we built a gene co-expression network from our scRNA-seq data using the algorithm implemented in *bigScale2*^46^, which uses a gene-gene correlation metric specifically tailored for sparse single-cell data. This resulted in a gene-gene network containing 5,238 nodes (genes) and 370,794 links (connecting two genes if their correlation was above 0.9). The biological validity of this network was confirmed by the fact that out of 59,545 genes both present in our network and in known TF-target lists (Arabidopsis Gene Regulatory Information Server, AGRIS^47^), 51,658 (∼86%) were preserved as linked pairs in our network. To identify groups of genes with correlated expression profiles, we used the Louvain algorithm and obtained a total of 16 gene modules (Fig S6, Supp Table 2), and summarised their expression as the first principal component of a PCA, which we refer to as an eigengene^48^. Among them, most of the modules were broad in all the trajectories with different temporal patterns. Module 6 seems to represent genes with high expression in PSE (Fig S6). In contrast, module 1, which contains 1,367 genes (Supp. Table 3), displays an increasing expression in both PPP and CC trajectories and a lower-than-average expression in PSE. Reporter lines for genes in this module followed these predictions: in addition to genes with broader expression (like *MES7*, Fig1a), we identified genes showing a ring pattern, expressed specifically in all the cells around PSE (Fig 3b, Fig S8). While *At3g11930*, *At2g02230* (*PHLOEM PROTEIN 2-B1*, *PP2-B1)*, *At5g47920*, *At3g16330*^2^ and its sister gene *At1g52140*, and *At4g27435* do not show expression in PSE, *At5g59090* (*SUBTILASE 4.12*, *SBT4.12), At2g20562* (*TAXIMIN 2*, *TAX2)* and *At1g26450* are expressed in late PSE in addition to being expressed in a ring pattern. Some of the genes found have broader expression (Fig S8). For instance, *At4g27435* is expressed in CC and occasionally in PPP and protoxylem plus lateral root cap. *At3g21770* (*PER30*) and *At3g11930* are found in the ring around PSE but extend to procambium higher up (Fig S8).

Out of nine genes with a ring expression pattern, seven were found in module 1, with *TAX2* (*At2g20562*) not included in our network and *At4g27435* found in module 4, which includes genes expressed in all trajectories. Despite module 1 being the largest on our network, this result is more than would be expected by chance (hypergeometric test, p-value = 0.0005).

Because of the large size of module 1, we tried to refine our analysis by sub-clustering the genes within this module, to identify a more specific group of candidate “ring genes”. This resulted in 15 sub-modules, with five of them containing over 100 genes (Fig S7). Six of the seven “ring genes” from module 1 fell within the same sub-module 1 (the exception was *At3g16330*), which again is more than would be expected by chance (hypergeometric test, p- value = 0.0009). While we do not expect that all of the 326 genes in this sub-module have a ring expression pattern, this analysis highlights that this pattern is widespread for a variety of phloem genes, which group together by similarity in expression pattern. On the other hand, a gene such as *MES7*, which we saw was not entirely ring-specific, fell in a different sub-module. Therefore, our network analysis suggests that there is a complex ring-specific pattern of expression shared across several genes in the phloem pole.

The complex patterns in the cells around PSE point out that PSE-adjacent cells share some common developmental programs that are maintained even when cells differentiate into their specific identities, suggesting the transcriptional signature of phloem cells is influenced by multiple positional cues.

This set of genes could be important to understand how PSE relates to its neighbouring cell types before and after enucleation. Indeed, as observed in the UMAPs, the ring pattern is frequent right after PSE enucleation, suggesting a shift in the phloem pole governance after PSE enucleation.

### *PINEAPPLE* genes extend the ring domain to the early phloem

Among the genes in module 1, sub-module 1, that also extend their expression into the most meristematic clusters, we found a DOF transcription factor, *DOF1.5* (*COGWHEEL1,* COG1). This gene and the sister gene *DOF2.3* (*CYCLING DOF 4*, *CDF4*), are expressed in early phloem cells (Fig 4a). *CDF4* is a differentiation factor in columella cells, repressed by WOX5^49^. The role of *COG1* in roots is unknown but it is a negative regulator of phytochrome signaling^50^ and promotes brassinosteroid biosynthesis by upregulating *PIF4* and *PIF5*, leading to hypocotyl elongation^51^. Both genes have been involved in regulating tolerance to seed deterioration^52, 53^ as well as flowering time^54^.

Transcriptional fusions of both genes confirmed the expression of both TF in PPP, CC and MSE from 40 µm from the QC, remarkably earlier than the other ring genes described above. While both genes form a ring around PSE reminiscent of a pineapple slice, *DOF1.5* (from now on *PINEAPPLE1*, *PAPL1*) is also expressed in the epidermis (Fig 4b) and *DOF2.3* (*PAPL2*) is found in columella cells with a broader domain towards the QC (Fig S9f). The ring pattern observed with the GFP fusion construct extends one layer towards procambium when fused to 3xYFP expression (Fig S9a), indicating a weaker expression in this layer. Translational fusions show these transcription factors are nuclear localised and not mobile (Fig 4d, Fig S9a- c, S9e), since transcriptional and translational patterns are coincident. This indicates that PAPL transcription factors act cell-autonomously. Together with the translational domain of *MAKR5*, the expression domain of *PAPL* genes indicate complex expression patterns in the phloem are relevant from an early stage.

*PAPL* genes, as other genes in module 1, were predicted to be *PEAR* targets^2^. PEAR transcription factors move to PSE-adjacent cells to control periclinal cell divisions and other transcriptional programs non-cell autonomously. This is evidenced by markers like *SAPL* and *At3g16330* becoming ectopically expressed after broad *PEAR* overexpression or *SAPL* being expressed in PSE upon PSE plasmodesmata closure^2^.

To validate *PAPL* genes as *PEAR* targets, *PAPL* reporter lines were expressed in *pear1pear2* double mutant, which resulted in a delay in *PAPL* expression, from 40 to 120 µm from the QC (Fig 4c, S9g). Since *PEAR* genes are highly redundant, we also introduced *PAPL1* constructs in the pear sextuple mutant, *pear sext*^33^, where we observed a loss in its usual meristematic expression (Fig S9d). In parallel, closing PSE plasmodesmata connections to the neighbouring cell types using *icals3m* tool, the ring expression of *PAPL1* is altered (Fig 4e) and overexpressing *PEAR1* leads to ectopic expression of *PAPL2* (Fig 4f). These results validate that *PAPL* genes are downstream of *PEAR* targets and indicate that PEARs are needed and sufficient to express *PAPL* genes in the early phloem.

In addition to the *PAPL* genes, we validated that some of the genes in module 1 act downstream of PEAR TF. Indeed, PEARs are sufficient to induce *SBT4.12*, *At3g11930*, *MES7* and *PER30*, since these genes become ectopically expressed upon induction of *PEAR2* expressed under a ubiquitous promoter (*pRPS5A*) (Fig S9i). In *pear sext*, the expression pattern of *PER30* and *MES7* was modified, while *SBT4.12* expression was decreased and *At3g11930* spread towards the meristem.

### *PAPL* genes link phloem morphogenesis and root physiology

Next, we decided to check if *PAPL* genes were downstream of *PEAR* genes to control periclinal cell divisions. Since *PAPL* expression is delayed in *pear1pear2* double mutant and absent from early phloem, we chose this mutant as a background to express *PAPL1* under the *WOODEN LEG* (*WOL*) promoter. When inducing *PAPL1* expression (20h treatment or germinated directly in beta estradiol and grown for 5 days), we did not observe a phenotype similar to *PEAR1* overexpression with increased periclinal cell divisions in the root^2^ (Fig S10a- h). A similar result was observed when *PAPL1* was overexpressed in the stele in wild type background (Fig S10i-p). These observations indicate *PAPL* genes do not control periclinal cell divisions downstream of *PEARs*.

To gain insight into the function of *PAPL* genes, and after checking *papl* single mutants didn’t show any obvious root phenotype, we generated double mutants (*papl1-1 papl2* and *papl1-2 papl2*). Bulk RNA sequencing identified *CYCLING DOF 2*, *CDF2*, as upregulated in *papl1- 1papl2* (supplementary table S4). This is another DOF transcription factor expressed in the cortex, pericycle and procambium, partially overlapping with *PAPL* expression (Fig S9h). Presuming this gene was upregulated to compensate for the lack of *PAPL* genes, we generated a triple mutant using a *cdf2* T-DNA allele^54^ (*papl1-1papl2cdf2, 3papl*).

The triple mutant root was shorter than wild type in several conditions (Fig S11a) but the effect was more pronounced growing the seedlings in media without sucrose (Fig 4g, Fig S11a,b). While wild type plants grown in media without sucrose often showed a bimodal distribution in terms of root growth (Fig S11a,b), the proportion of roots arresting growth in 3*papl* was higher (Fig 4g, Fig S11e). Even if there is high variation between seed batches, the mutant root seems to be on average lower than the average root length in wild type (Fig S11d). Contrary to other phloem development mutants, adding 1% sucrose to the media mostly suppressed the mutant phenotype (Fig 4h).

Since the mutants could be rescued by transferring them to sucrose, we aimed to identify the time point at which sucrose is needed for 3*papl*. For this experiment, we transferred plants from sucrose supplemented to sucrose-depleted media and *vice versa*. The more time the mutant seedlings spent without sucrose (blue series), the more difficult it was for them to recover root growth (Fig 4h, Fig S11g). Those recovering managed to grow well (Fig S11e). Spending at least 3 days in sucrose (red series) was necessary for the mutant seedlings to grow normally while spending only two days in sucrose was not enough for root growth recovery (Fig 4h, Fig S11g). In the confocal, the root meristem of seedlings that got arrested, looked shorter and stunted (Fig S11c). *PAPL* genes were expressed at this stage in both sucrose and non-sucrose conditions showing similar patterns as observed in more mature seedlings and phloem unloading marker *pSUC2::GFP* showed no phenotype in the mutant background in any sucrose content (Fig S11f).

To better understand the *3papl* phenotype, we carried out metabolic profiling of leaves and roots of seedlings grown in a sucrose-depleted media across six developmental stages (2-7 days post-sowing, dps) (Supplementary Table S5). We identified 7 and 5 metabolites in leaves and roots, respectively, with significant differences between WT and mutant in at least one of the time points (<5% false-discovery rate from a linear mixed model fit to the whole data, see methods; Fig S12a). One of those metabolites was sucrose, with a significant difference only in the roots, where it started at lower levels in the mutant (days 2 and 3) and then continued to increase to reach levels comparable to the WT at the end of the experiment at day 7 (Fig S12b). A similar pattern, with more significant points, was observed in fructose, which is a component of sucrose, and to a lesser extent in glucose, the other monosaccharide forming sucrose (Fig S12b). It has been described that by the time the radicle emerges, all the sugars stored in the Arabidopsis seed have been consumed. Within 48 hours after germination, lipid and protein reserves are exhausted and seedlings need to switch to autotrophic growth ^55, 56^. The data suggest *PAPL* genes could be important after the seedling has transitioned to autotrophic growth, facilitating sugar transport to sink tissues like roots. The continued increase in sucrose in the mutants could be due to the, on average, smaller size of *3papl* seedlings and stunted growth, which could therefore lead to reduced sucrose consumption and therefore its observed continued accumulation.

## DISCUSSION

Our manuscript demonstrates the power of tissue-specific transcriptomes combining FACS and single cell sequencing to study elusive cell populations underrepresented in organ general cell atlases. The use of droplet-based technologies also allowed us to gather more cells and a higher resolution than plate-associated methods.

The phloem pole cell atlas is allowing a holistic understanding of phloem. While there are specific genes for PPP and CC, these cell types share the first stages of their developmental trajectory. Trajectory analysis also revealed the connection between CC and early MSE, providing new insights on early stages of MSE development. The commonalities among the different cell types were validated by gene regulatory network analysis and reporter lines confirmed the relevance of the ring expression pattern in all the cells around PSE.

PSE differentiation involves enucleation and becoming dependent on adjacent cells for survival. Using *APL* expression as a standard, we mapped the enucleation point in the atlas. While PSE organizes the phloem pole in the meristem neighboured by unspecialized cells, PSE enucleation marks the onset of cell differentiation for adjacent cells and switches on similar gene regulatory networks in PSE-surrounding lineages, as evidenced by the ring pattern shown by many genes right after PSE enucleation.

The coordinated expression in the cells of the phloem pole highlights the importance of positional information and cell to cell communication to preserve phloem function when PSE delegates control in the adjacent cells. They also underpin the relevance of PPP cells, which we believe should be considered a built-in part of phloem.

A phloem plasticity zone was recently described in the root meristem, when CC and MSE cells could act as a reservoir for PSE identity^17^. This further supports the coordination between the pole identities to ensure correct phloem morphogenesis. It would be interesting to investigate if PPP can also transdifferentiate to PSE if required.

In turn, the similarities between root pericycle cells and phloem parenchyma cells in leaves suggests parenchymatic cells share characteristics despite being present in different organs with variable anatomic configurations and reinforces PSE as the phloem pole organiser.

As evidenced by *PAPL* expression patterns, the cells around PSE share transcriptional characteristics from an early stage and seem to be important in the transition to autotrophy. Contrary to other phloem morphogenesis mutants, like *apl*, the presence of sucrose in the media almost completely suppresses the root growth phenotype of *3papl*. Since phloem is in charge of nutrient transport and a smaller amount of sucrose and its component fructose is detected in both mutant leaves and roots at 3 dps when root anatomy is comparable between wt and mutant, we interpret *PAPL* genes regulate nutrient allocation between the leaf source organs and the root sink in young seedlings, when embryo reserves are scarce. *PAPL* genes could either regulate phloem loading, long distance transport or phloem function and more studies are required to determine their precise function.

## MATERIALS AND METHODS

### Plant growth conditions

All *Arabidopsis thaliana* lines used in this study were in Col-0 background except *pear1* mutant allele, which is in Ler background, conferring *pear1pear2* mutant a mixed Ler appearance. Plants were grown in ½ MS Basal salts media (0.5 MS Salts, 1% Difco agar, with or without 1% sucrose) at 23°C and long day conditions, except for sorting experiments, when they were grown using 1x MS Basal salts at 23°C with 30% humidity and 188 μM of light, long day conditions, to be able to compare with other transcriptomic data.

*papl1-1* (*cog1-6*, from gene *At1g29160, PAPL1, DOF1.5, COG1)* has a single nucleotide deletion (G) at position +85, which generates a premature stop codon. This mutant was identified as a *cog1-D* suppressor^51^. *papl2* (*At2g34140, PAPL2, DOF2.3, CDF4)* has a 4 bp deletion (CAAG) at position +99 creating a premature stop codon. The *cdf2* T-DNA allele (GK782H09) is a knockdown allele^54^. Triple mutant was obtained by crossing the double mutant *papl1-1papl2* to *cdf1r235*^54^, selecting for mutant *3papl* and homozygous wild type alleles for all other genes.

5 µM Beta estradiol or 10 µM DEX were used in the inducible constructs for the indicated times. Plants induced with DEX were treated for 24 hours.

### Sorting and single cell sequencing

Seedlings from the different marker lines were grown vertically over mesh (Normesh, 100 µm) for five days in the conditions specified above. Approximately one third of the root including the root tip was chopped with razor blades and the tissue transferred to the protoplasting solution for an hour as in ^55^. In the case of the sample “MAKR5 enriched in root tips”, we submerged the root tips of intact roots in eppendorfs containing the protoplasting solution for 15 minutes, which is enough time for the meristems to be enzymatically cut from the rest of the root. Then the separated root tips were incubated for 45 minutes and treated as the other samples. Washed protoplasts suspended in solution A were taken at room temperature to the sorting facilities. For the gating, a wild type Col0 sample was run first to establish the fluorescent negative gate. Then this sample was subsequently stained with DAPI and DRAQ5 to gate for intact cells that contained DNA, respectively. The corresponding sample containing fluorescent protoplasts was then stained subsequently with DAPI and DRAQ5 and underwent FACS. Gating helped enrich intact (DAPI negative), YFP/GFP positive, DNA containing cells (DRAQ5 positive) that were sorted with a 130 µm nozzle using a High speed Influx Cell Sorter (BD Biosciences). Sorted protoplasts were harvested in W5 solution (154 mM NaCl, 125 mM CaCl 2, 5 mM KCl, 5 mM MES (2-(N morpholino)ethanesulfonic acid) in BSA coated 1.5 ml Eppendorf tubes. Cells were centrifuged for 12 minutes at 200g to eliminate the excess of supernatant. Immediately, Single-cell RNA-seq libraries were prepared in the Cancer Research UK Cambridge Institute Genomics Core Facility using the following: Chromium Single Cell 3′ Library & Gel Bead Kit v3, Chromium Chip B Kit and Chromium Single Cell 3’ Reagent Kits v3 User Guide (Manual Part CG000183 Rev C; 10X Genomics). Cell suspensions were loaded on the Chromium instrument with the expectation of collecting gel-beads emulsions containing single cells. RNA from the barcoded cells for each sample was subsequently reverse-transcribed in a C1000 Touch Thermal cycler (Bio-Rad) and all subsequent steps to generate single-cell libraries were performed according to the manufacturer’s protocol with no modifications. cDNA quality and quantity was measured with Agilent TapeStation 4200 (High Sensitivity 5000 ScreenTape) after which 25% of material was used for gene expression library preparation.

Library quality was confirmed with Agilent TapeStation 4200 (High Sensitivity D1000 ScreenTape to evaluate library sizes) and Qubit 4.0 Fluorometer (ThermoFisher Qubit™ dsDNA HS Assay Kit to evaluate dsDNA quantity). Each sample was normalized and pooled in equal molar concentration. To confirm concentration pool was qPCRed using KAPA Library Quantification Kit on QuantStudio 6 Flex before sequencing. All samples were sequenced using Illumina NovaSeq6000 sequencer with following parameters: 28 bp, read 1; 8 bp, i7 index; and 91 bp, read 2.

### Analysis of single-cell RNA-seq

Here we give a briefer description and overview of our analysis steps, but the full details of our analysis pipeline (e.g. specific package functions and options used) can be seen in our code repository at https://github.com/tavareshugo/publication_Otero2021_PhloemPoleAtlas.

To obtain unique molecular identifier (UMI) counts for each gene, the raw sequencing reads were aligned to the reference Arabidopsis TAIR10 genome using the Araport11 gene annotation (both downloaded from Ensembl release 45) using 10x Genomics Cell Ranger v3.1.0^56^. The data were processed and quality-filtered using several Bioconductor packages^57^. Empty droplets were inferred and removed using dropletutils v1.8.0^58^, and data normalisation was done using both the pooling method implemented in scran v1.16.0^59^ and the variance-stabilising transformation from sctransform v0.2^34^. To adjust for potential batch effects, data from the different samples (i.e. sorted with different GFP fusion markers and/or from different public datasets) were integrated using the Mutual Nearest Neighbours (MNN) algorithm implemented in batchelor v1.4.0^35^. After initial data exploration and quality checks, we retained cells with at least 2000 detected genes and genes detected in at least 100 cells (a gene was considered to be detected if it had at least 1 UMI count). Downstream analysis was done on these filtered data, batch-normalised using MNN and using variance-stabilised transformed values. However, our conclusions were qualitatively robust to the specific choice of normalisation methods.

Cell clustering was performed by first defining a “shared nearest neighbours” graph and then identifying modules in the graph using the Louvain algorithm (using scran v1.16.0^59^ and igraph v1.2.6^60^. To annotate our cells we used a set of genes with known expression patterns (from promoter fusion microscopy experiments) and calculated, for each cluster, the percentage of cells where each marker gene was detected as well as the (z-score scaled) average expression of the gene in that cluster.

The same pipeline was applied to the public datasets, also integrated using MNN. The quality of this data integration was confirmed by checking that the majority of our annotated cells were clustering together with the same cell types in other datasets. We produced two sets of data integration, one with root data and another with leaf data. Details of the public datasets used are given in (Supplementary Table 6).

To explore how well cells from leaf and root datasets mixed in clusters where they co-occurred (namely a cluster containing both leaf phloem parenchyma and root phloem pole pericycle cells) we used the shared-nearest-neighbours cell graph used for clustering and calculated the proportion of edges between root-leaf cells (the vertices of the graph). This value was then compared with a null expectation, obtained by randomly shuffling the cell tissue labels 1000 times and calculating this proportion each time. The 95% inter-percentile range of this null distribution was then used to compare with the observed value.

To further temporally annotate our phloem pole atlas dataset we used several approaches. Early dividing cells were identified by checking the expression of all annotated cyclins and other cell cycle markers such as *AUR1* (*AT4G32830*) and *KNOLLE* (*AT1G08560*). We also cross-referenced our data with a published dataset that profiled the transcriptome of longitudinal root sections using microarray technology^1^. Based on 9,674 common genes between the two datasets, we assigned each of our cells to the longitudinal section of Brady et al. that had the highest Spearman correlation with it. We also used the RNA velocity method implemented in scVelo v0.2.2 to infer developmental dynamics in our data^42^. Finally, cells were assigned to lineages and ordered by pseudotime using slingshot v1.6.0^41^. In this latter case we used a semi-supervised approach, where the starting point for the inferred trajectories was set to the cluster highly expressing cell-cycle markers and identified as the earliest cluster when cross-referencing with the Brady et al. dataset. In this manner we obtained biologically meaningful trajectories (without setting this constraint several more trajectories were obtained but with an ordering of cells which was the reverse of what was expected from our other analyses). We obtained smooth gene expression patterns for each trajectory using generalised additive models, as implemented in tradeSeq v1.2.0, which were then used to explore gene expression patterns along the slingshot trajectories.

To cluster genes based on their similarity of expression across the cells, we built a co-expression network using a modified version of bigSCale^46^, adapted to work on any species (rather than the original version suited only for mouse and human). The modified package is available from https://github.com/tavareshugo/bigSCale2/tree/support-any-species.

Summarily, bigScale builds a gene correlation matrix not from the original count data (which in scRNA-seq is too noisy and sparse), but from a z-score statistic calculated between pairs of cell clusters. These clusters are iteratively generated to ensure the z-scores capture as much diversity in gene expression patterns across the cells as possible. In this way, correlations between genes are more robust to the noisy and sparse nature of single-cell RNA-seq data. This correlation matrix was then thresholded at 0.9 to obtain a gene-by-gene adjacency matrix, resulting in a network with 5,238 nodes (genes) and 370,794 links. We identified gene modules using the Louvain algorithm, resulting in 16 modules. From each module, we calculated an eigengene following the procedure in WGCNA vX^48^, which essentially summarises the expression of all genes of a module as the first principal component score from a principal components analysis (PCA) done on those genes. The largest of these modules - module 1 containing 1,367 genes - contained several genes of interest for our analysis, and was therefore re-clustered with Louvain to generate 15 sub- modules. This was further justified by the fact that the variance explained by this module’s eigengene was relatively low (21.44%), suggesting some heterogeneity in expression patterns within the module. To further interpret these results, the eigengenes from these sub-modules were joined with the pseudotime trajectories from slingshot, although we note that no information about trajectories was used to build the network itself. Therefore, the fact that the different approaches (gene network and pseudotime analysis) reveal groups of genes with similar patterns of expression is a strengthening point in our analysis.

### Generation of reporter lines and confocal images

*Promoter::VENUSer* fusions were generated for the genes *At3g27030*, *At2g23560* (*MES7), At2g32210, At5g64240 (MC3), At5g58690* (*PLC5*), *At2g38640*, *At3g11930*, *At2g02230* (*PP2- B1)*, *At5g47920*, *At4g27435*, *At5g59090* (*SBT4.12), At3g21770* (*PER30*), *At1g26450, At1g29160 (PAPL1, DOF1.5, COG1), At2g34140 (PAPL2, DOF2.3, CDF4), At5g39660 (DOF5.2, CDF2)*.

Translational fusions were also generated for *At2g20562* (*TAX2), PAPL1* and *PAPL2*. 3xYFP constructs were also generated for transcriptional fusions of *PAPL1* and translational fusions of *PAPL1* and *MAKR5*.

Promoter fragments between 622-4879 bp were amplified by PCR and cloned using MultiSite-Gateway (Supplementary Table 7). Transcriptional fusions to *VENUS* with an ER tag or translational fusions to YFP were generated in vectors with either resistance to Basta or Hygromycin or a Fast Green/Fast Red selection system. All the constructs were transformed in Col0 background and at least 2 independent lines were analysed for each.

Roots from 5-7-day-old seedlings were either imaged in the confocal directly after mounting them in 50 µg/ml propidium iodide or fixed for 45 minutes in 4% paraformaldehyde in PBS and cleared using ClearSee solution as in^61^. Cleared roots were then stained with SCRI Renaissance 2200 and observed under the confocal. Images were acquired at 512x512 resolution using the confocal Leica SP8.

### Bulk RNA-seq transcriptomics

Wild type and *papl1-1papl2* seedlings were grown on mesh in ½ MS media with sucrose in the above-mentioned conditions for 5 days. Root meristems from wild type and mutant were manually and individually dissected in parallel under a stereomicroscope using 18G needles. Meristems were preserved in RNAlater RNA stabilization reagent (Qiagen) until 120 meristems per replicate were gathered. 4 replicates for each mutant and wild type were used for RNA extraction.

RNA was extracted using the RNeasy Plant Mini kit from Qiagen and RNA integrity and concentration were checked using TapeStation and Qubit 2.0 fluorometer (Life Technologies) respectively. After quality control in Novogene company, the best 3 replicates for mutant and wild type were used for library construction and sequencing following the Novogene pipeline. Briefly, mRNA was enriched by using oligo dT beads and fragmented randomly. cDNA synthesis was performed using random hexamers and reverse transcriptase. After first-strand synthesis, the second strand is synthesized by nick-translation. Library is ready after a round of purification, terminal repair, A-tailing, ligation of sequencing adapters, size selection and PCR enrichment. Library concentration was quantified using a Qubit 2.0 fluorometer (Life Technologies), Insert size was checked on Agilent 2100 and quantified more accurately by quantitative PCR. Libraries were fed into the HiSeq XTEN platform for sequencing. Original raw data were transformed to Sequence Reads by base calling and raw data recorded in FastQ files. Low quality reads or reads containing adaptors were filtered out. TopHat2^62^ v2.0.12 was used to map the reads to the reference genome (TAIR10). HTSeq^63^ v0.6.1 software was used to analyze the gene expression level using the union mode. Fragments Per Kilobase of transcript sequence per Millions base pair sequenced (FPKM) value of 0.1 or 1 was set as the threshold to determine whether a gene is expressed or not. To compare gene expression levels under different conditions, FPKM distribution diagram and violin plot were used. For biological replicates, the final FPKM would be the mean value. The differential gene expression analysis consisted of read-count normalization, model-dependent mean value estimation and FDR value estimation based on multiple hypotheses testing. DESeq^64^ v1.10.1 software was used for these steps.

### Metabolic profiling

Arabidopsis plants were grown across six developmental stages (from day to day 7) on mesh in solid media containing sucrose or devoid of sucrose. Each day of the time course, leaves and roots were harvested separately and snapped frozen in liquid nitrogen. 50 mg of leaves and 20 mg of roots were ground using a Tissue Lyser. Extraction was performed according to Lisec et al. (2006)^65^, with modifications. In detail, 750 µl/300 µl of extraction buffer (100% methanol plus the internal standard adonitol, Sigma) were added to root and leaf samples respectively. Samples were vortexed and transferred to a shaker at 70 ℃ for 15 minutes. 375 µl/200 µl of chloroform and 750 µl/350 µl of water were added to the tubes for leaves and roots respectively, and samples were centrifuged for 10 minutes at maximum speed. 400 µl (roots) and 200 µl (leaves) of supernatant were dried for each sample using the speedvac. Samples were kept at -80℃ until processing.

The dried samples were derivatized for 2 hours at 37 °C in 50 `μ`I of 20 mg ml-1 methoxyamine hydrochloride (Sigma-Aldrich, cat. no. 593-56-6) in pyridine followed by a 30 min treatment at 37 °C with 100 `μ`I of N-methyl-N-(trimethylsilyl)trifluoroacetamide (MSTFA reagent; Macherey-Nagel, cat. no. 24589-78-4). For each sample, 1 `μ`I was injected in splitless mode to a chromatograph coupled to a time-of-flight mass spectrometer system (Leco Pegasus HT TOFMS; Leco Corp., St Joseph, Ml, USA), using an autosampler Gerstel Multi-Purpose system (Gerstel GmbH & Co.KG, Mulheim an der Ruhr, Germany). Chromatograms and mass spectra evaluation, as well primary metabolites identification based on the expected retention time and mass fragmentation were performed using the software Xcalibur software (Thermo Fisher Scientific).

To estimate differences between WT and *3papl* metabolite levels, we fit a hierarchical model to the data adding terms for genotype, stage (dps), tissue and their interactions. We included a random effect to account for multiple measurements per sample (each sample contributed 21 data points, one for each metabolite). Due to the skewed distribution of peak areas, the data were modelled on a log-scale, which produced well-behaved normally distributed residuals. From this model, we obtained estimates of the difference between the two genotypes for each metabolite and tissue, using the *emmeans* v1.6.2-1 R package. The p-values from the *emmeans* contrasts were corrected for multiple testing using the false discovery rate method.

Additional information on metabolomics analysis and metabolites annotation are reported in Supplementary table 8 (sheets checklist and overview) according to the guidelines provided in Alseekh et al^66^.

## Data availability statement

Sequencing data from 10x Chromium single-cell RNA-seq is available from NCBI’s Gene Expression Omnibus through GEO accession number GSE181999: https://www.ncbi.nlm.nih.gov/geo/query/acc.cgi?acc=GSE181999.

Sequencing data from bulk RNA-seq is available from NCBI’s GEO accession number GSE182672: https://www.ncbi.nlm.nih.gov/geo/query/acc.cgi?acc=GSE182672.

All other data (phenotypic scoring, microscopy imaging, plasmid maps) are available from the Cambridge Apollo Repository (https://doi.org/10.17863/CAM.74836).

A persistent DOI will be available upon acceptance.

Analysis code, with instructions on how to run it, is available from: https://github.com/tavareshugo/publication_Otero2021_PhloemPoleAtlas.

## Acknowledgements

We thank Wolf Frommer and Ji-Yun Kim for providing *pSWEET11:SWEET11-2A-GFP* seeds, Christian Hardtke for providing *pMAKR5:MAKR5-GFP* seeds and plasmids, Chiara Cossetti, Reiner Schulte and all the staff from Flow Cytometry Core Facility at CIMR for their technical support with cell sorting, Katarzyna Kania from Genomics Unit at CRUK for preparing Single-cell RNA-seq libraries, Bruno Guillotin and Kenneth Birnbaum for helpful insights on single cell analysis, Raymond Wightman and Gareth Evans for technical support with microscopy experiments, Gill Hindle, Jemma Salmon and Sally Ward for media preparation, Karolina Blajecka for technical assistance, Kristina Petkovic and Rut Alcaina for technical support. S.O. was supported by a Herchel Smith postdoctoral fellowship from the University of Cambridge (2017-2020). L.K. received funding from the SNSF (P2LAP3_178062) and a Marie Curie IEF (No. 795250). This work was supported by Finnish CoE in Molecular Biology of Primary Producers (Academy of Finland CoE program 2014-2019) decision #271832, the Gatsby Foundation (GAT3395/PR3)), University of Helsinki (award 799992091) and the ERC Advanced Investigator Grant SYMDEV (No. 323052). A.R.F and V.D. acknowledge support from the Max-Planck Society.

## Author Contributions

S.O. performed the experiments, I.S. identified *PAPL1* and *At3g16330* expression patterns which appeared as *PEAR* targets in microarray data, P.R. provided *pear1pear2* double mutant, *pPEAR(del)::3xYFP* and advised on experimental design, Y.L. and H.T. analysed gene regulatory networks, P.R., M.B., L.K.,B.B. and J-o.H. participated in sample collection for sorting and metabolomic profiling, J-o.H. provided *pSAPL::YFPer* line, V.D. and A.R.F. carried out the metabolic profiling and data analysis, F.P. and T. L. provided the *papl2* and *papl1-2* alleles, H.T. designed and performed the single cell data and statistical analysis, S.O., H.T. and Y.H. conceptualised and designed the study. S.O. wrote the manuscript with input from Y.H., H.T., P.R. and L.K. All authors read, edited and discussed the manuscript.

## Competing Interests statement

The authors declare no competing interests.

---

**Fig S1.**
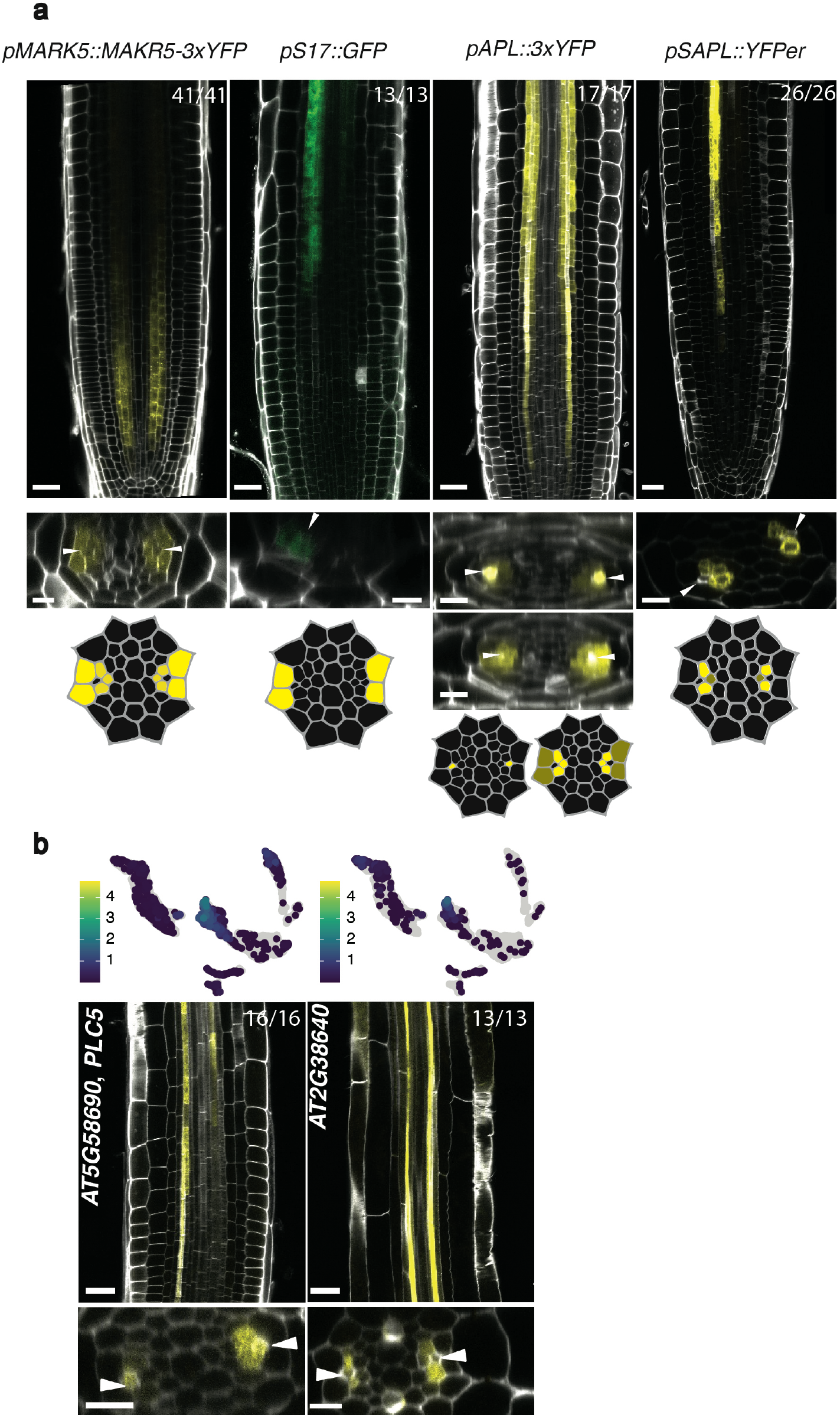
Markers used for sorting phloem pole cells and new CC genes identified. a) *MAKR5* translational fusion highlights all the cells surrounding PSE^1^ from the quiescent centre (QC) (often stronger from cell number 3) until the differentiation zone, where it becomes weaker. In mature parts of the root this marker spreads to the whole pericycle. Published marker was fused to 3xYFP to increase signal. S17^2^ is expressed in PPP from the unloading zone. *pAPL::3xYFP* is expressed first in PSE and after enucleation switches to all the cells around PSE, stronger in CC. This line is not fully reflecting *ALTERED PHLOEM DEVELOPMENT* (*APL)* endogenous expression, since it has some expression in the outer layers but it is a very strong phloem pole marker. *sAPL* is expressed in CC and MSE (weaker) from 90-120 µm from the QC. *pPEAR1(del)::3xYFP* is a modified version of the *PEAR1* promoter that is expressed in early PSE, MSE and a procambial cell resulting from the same division plus columella cells. See Roszak et al. ^3^ for the detailed expression pattern b) Newly identified genes expressed in CC (*PLC5*, *At5g58690*, and *At2g38640*). UMAPs show the particular cluster-weighted normalised expression of each gene in the phloem pole cell atlas and microscopy pictures are representative images of the transcriptional reporter lines where the gene promoter is fused to *VENUSer*. Scale bar in the longitudinal sections is 25 µm while it is 10 µm in the cross sections. White arrowheads point to PSE cells as a reference point. The numbers in each panel indicate samples with similar results, of the total independent biological samples observed.

**Fig S2.**
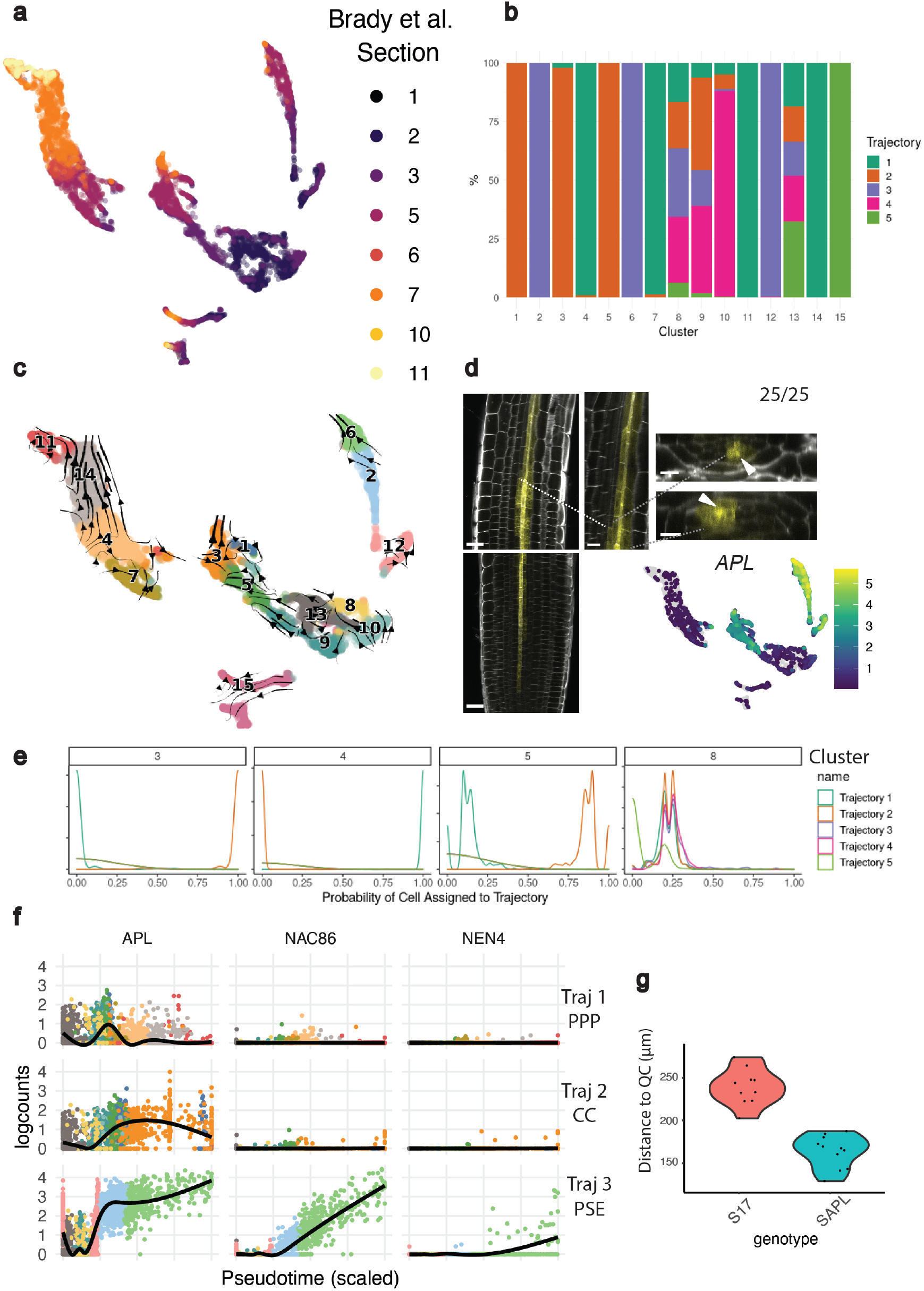
Validation of the temporal information in the UMAP by using complementary tools. a) Color-coded UMAP according to the longitudinal sections in Brady et al. 2007, showing a developmental progression in each cell lineage. b) Bar plot indicating the percentage of cells contributed by each cluster to each of the Slingshot trajectories shown in the main text. Bars are coloured by trajectory. c) RNA velocity analysis using scVelo, with velocity vectors projected on our UMAP d) Confocal pictures of *pAPL::VENUSer* showing continuous expression in PSE in the early root meristem with a patchy expression in the neighbouring cells that gets stable in CC and MSE shootward. As observed in the zoom picture, the signal in PPP gets weaker after PSE enucleation. The pictures are accompanied by a UMAP showing *APL* cluster-weighted normalised expression in the phloem pole cell atlas. Scale bar in the longitudinal sections is 25 µm while it is 10 µm in the cross sections and zoom. White arrowheads point to PSE cells as a reference point. 25 independent seedlings from three different lines expressing this construct were observed e) Probability of a cell to be assigned to different trajectories, ranging from 0 to 1. In the image we are showing a few clusters as an example. f) The expression of *APL* and PSE enucleation markers *NAC086* and *NEN4* was plotted along the PPP, CC and PSE trajectories, with the cells coloured by cluster number in the UMAP. *NAC086*, an *APL* target, appears later than *APL* in the PSE trajectory, and *NEN4*, a *NAC086* target, appears later than *NAC086*, indicating our approach matches the temporal aspects observed in PSE biology in the roots. g) Initiation of *S17* and *SAPL* gene expression. Distance from QC (µm) was measured to the first cell expressing each gene in 7 days post sowing (dps) seedlings, n=10 for *S17* and n=11 for *SAPL*.

**Fig S3.**
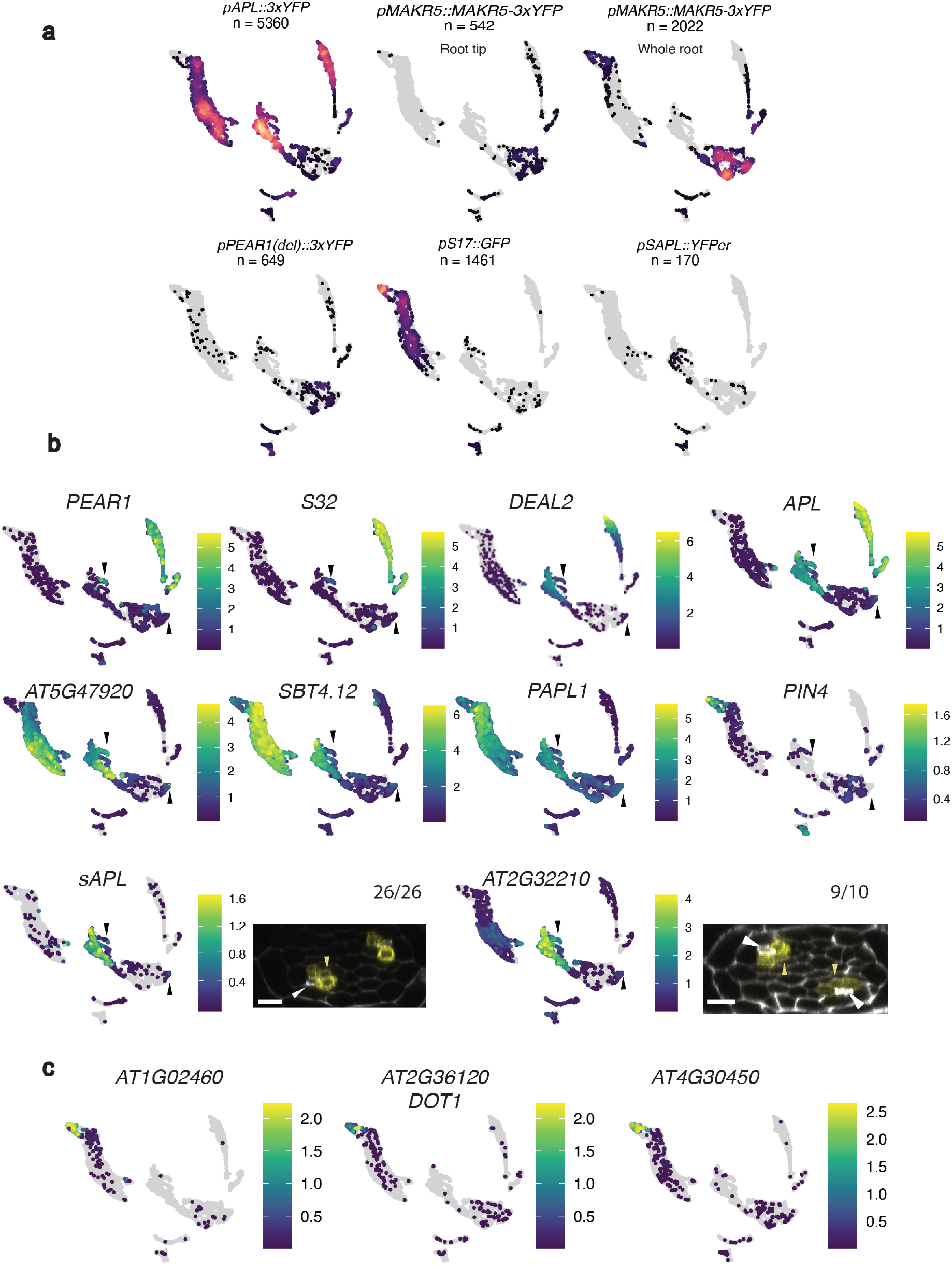
MSE cells identification and identity of cluster 11. a) Cells were plotted in the UMAP separated by sorting experiment, as indicated in each panel, to show which sorting experiment provided every cell. Colour indicates point density (lighter colour indicates higher density of points), with grey areas meaning an absence of cells. Numbers in each panel indicate the number of filtered cells contributed by that sorting experiment. We sorted *MAKR5* twice, one enriching in root tips (*MAKR5*) and another sorting the usual one third of the root (*MAKR5* differentiated). b) UMAPs showing the cluster-weighted normalised expression of marker genes used to identify MSE identity. *PEAR1*, *S32* and *DEAL2* are expressed in sieve elements. *APL* is genuinely expressed in PSE, CC and MSE. *At5g47920*, *SBT4*.*12* and *PAPL1* are expressed in a ring expression pattern, including MSE and other cell types. *PIN4* is used as a negative control, since it is excluded from sieve elements early in development^3^. *SAPL* and *At2g32210* are expressed in CC and MSE. Black arrowheads point to clusters 1 and 10. In the confocal cross sections, the scale is 10 µm. White arrowheads point to PSE as a reference point and yellow arrowheads point to MSE. The numbers over each picture indicate samples with similar results, of the total independent biological samples observed c) UMAPs for xylem pole pericycle markers, which are found in cluster 11 together with other PPP markers, indicating this is a late pericycle cluster.

**Fig S4.**
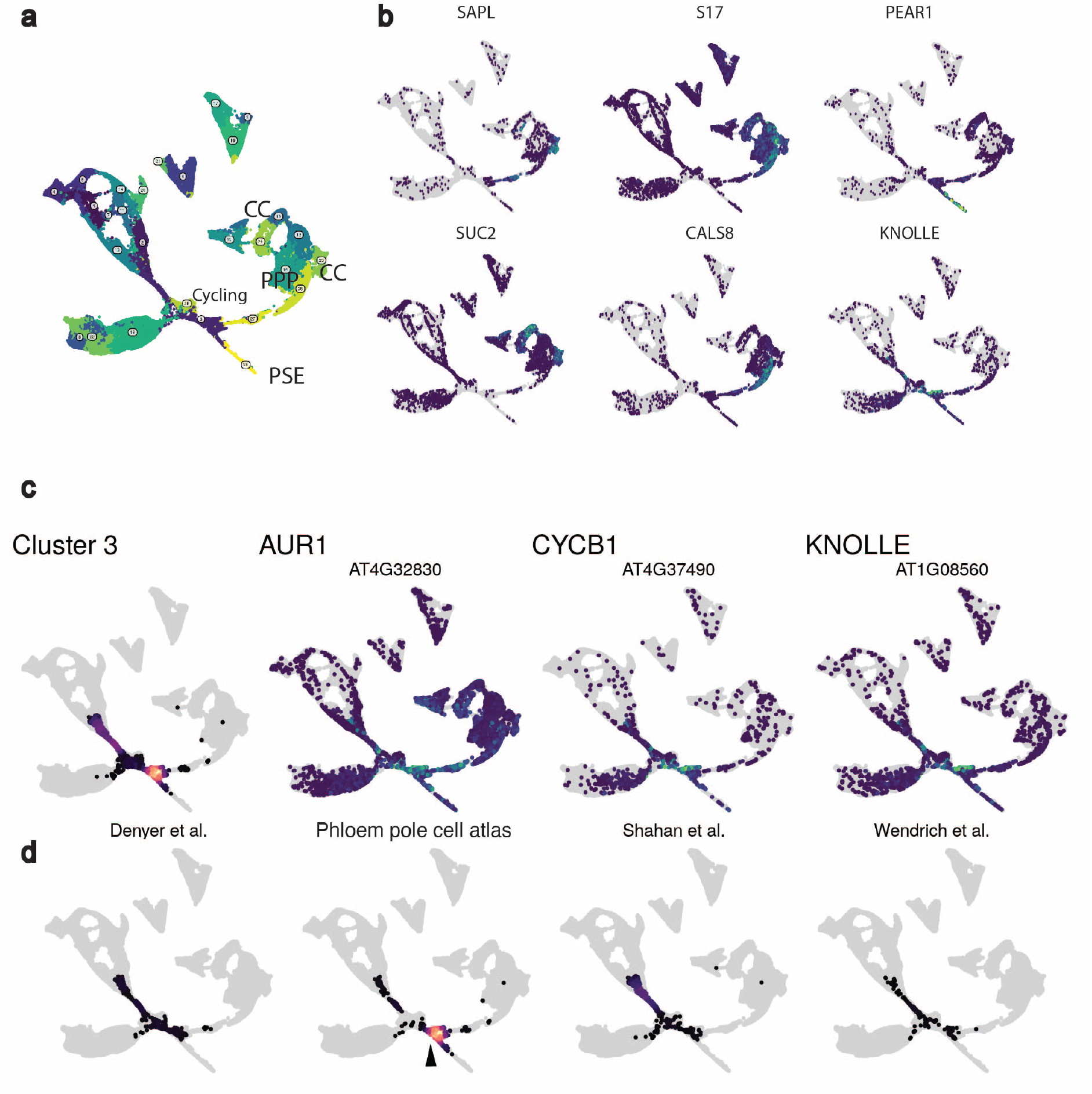
Phloem cell types in the integrated UMAP. a) A new UMAP containing 113340 cells was generated by integrating cells from Denyer et al. 2019^4^, Wendrich et al. 2020^5^ and Shahan et al. 2020^6^. Colours are used to differentiate cell clusters. b) Different markers were plotted in the UMAP to identify the phloem pole cell types: *SAPL* (CC and MSE), *S17* (PPP), *PEAR1* (PSE, MSE), *SUC2* (mature CC), *CALS8* (PPP and CC), *KNOLLE* (cycling cells). c) Cluster 3 of the integrated dataset containing root early cells and dividing cells. Cells in cluster 3 (early cells) are indicated in the first panel while the other panels in the row show the expression of G2/M genes in the integrated dataset, marking dividing cells d) Contribution of each single cell project to cluster 3. Observe the grouping of early phloem cells (black arrowhead) compared to the higher dispersion of early cells in other datasets.

**Fig S5.**
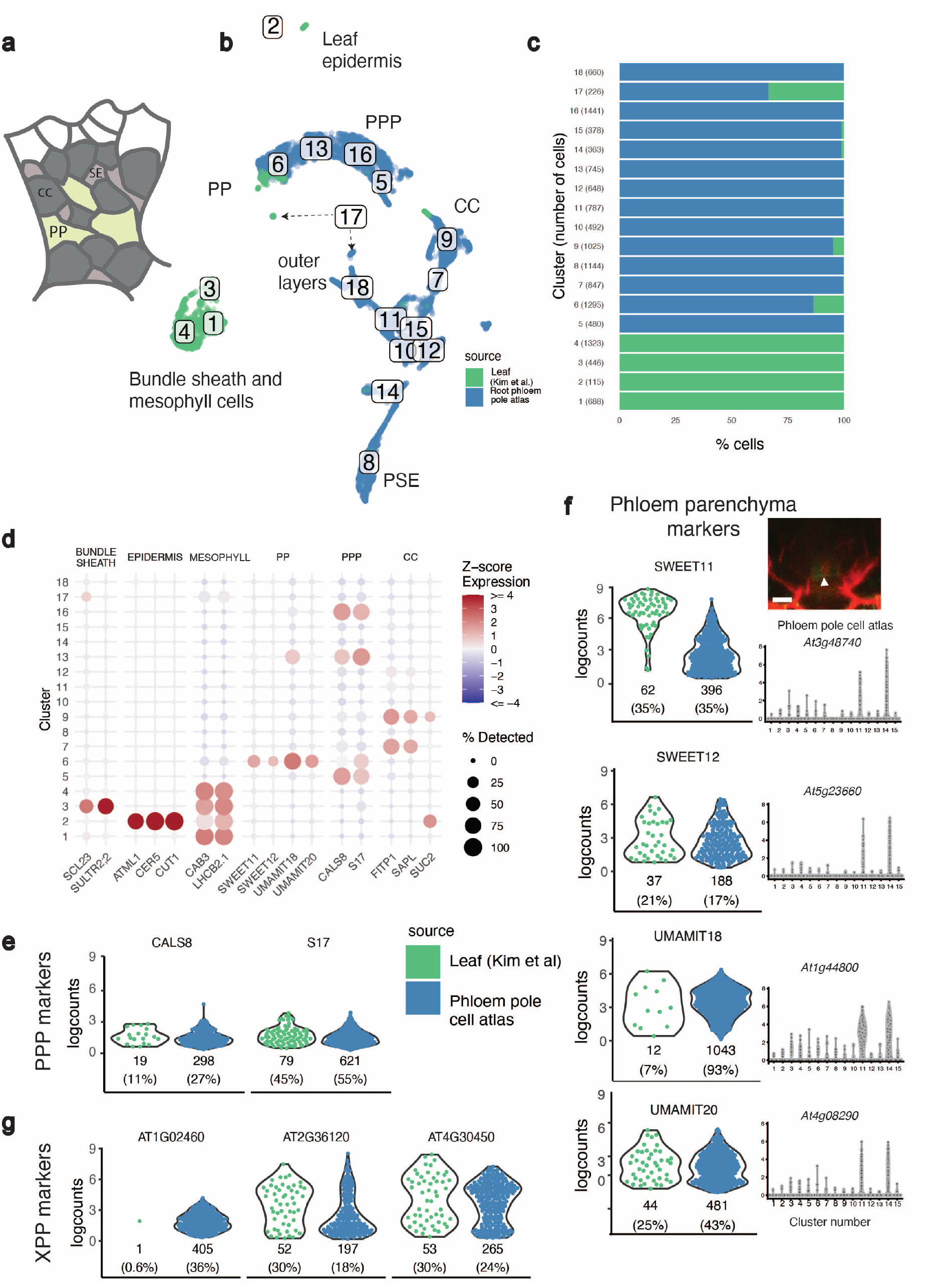
Integration with leaf single cell data reveals similarities in the gene expression between leaf phloem parenchyma and root pericycle. a) Schematic of the leaf minor vein showing phloem anatomy. Notice the different composition in terms of cell identities, cell number and organization compared to the root. Adapted from^46^ b) UMAP integrating the phloem pole cell atlas with the leaf single cell dataset. Cells were combined and reclustered, coloured by source (leaf in green, root in blue). Notice the separation of the leaf specific clusters (bundle sheath and mesophyll cells) and overlap in clusters 6 (PPP / phloem parenchyma) and 9 (CC). c) Percentage of cells contributed by each dataset in each cluster. Y axis shows the cluster number with the number of cells in it between brackets. Root cells are coloured in blue, leaf cells are in green. d) Cluster annotation of the root-leaf UMAP based on markers with known tissue-specific expression. The size of the points represents the percentage of cells in a cluster where the gene was detected (i.e. at least 1 UMI). The colour shows the scaled average expression of the gene (z-score, i.e. number of standard deviations above/below the gene’s mean across all cells) e) Violin plots showing the expression of PPP markers in leaf (green) and root (blue) cells for phloem pole pericycle markers (e), phloem parenchyma markers (f) and XPP markers (g) The confocal picture in f shows the expression of *pSWEET11::SWEET11-2A-GFP* in PPP in roots. In f, the gene expression of the respective genes is shown in the phloem pole cell atlas. The numbers under the blue/green violin plots indicate the number of cells in cluster 6 of the leaf/root UMAP expressing each gene, with the percentage between brackets. In the confocal picture, the scale is 10 um and the white arrowhead points to PSE as a reference point.

**Fig S6.**
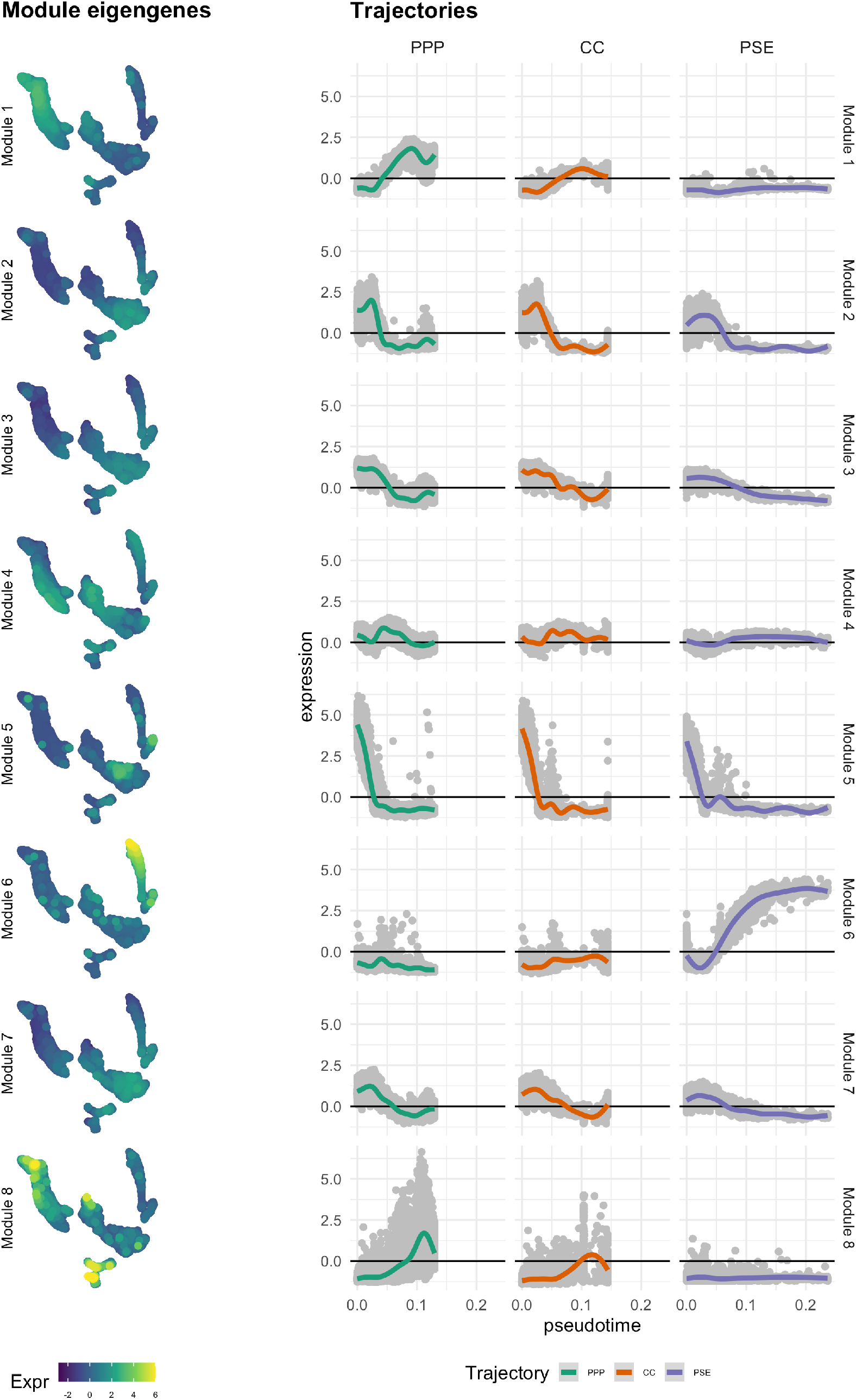
Part I. See next page for Part II and caption.

**Fig S6.**
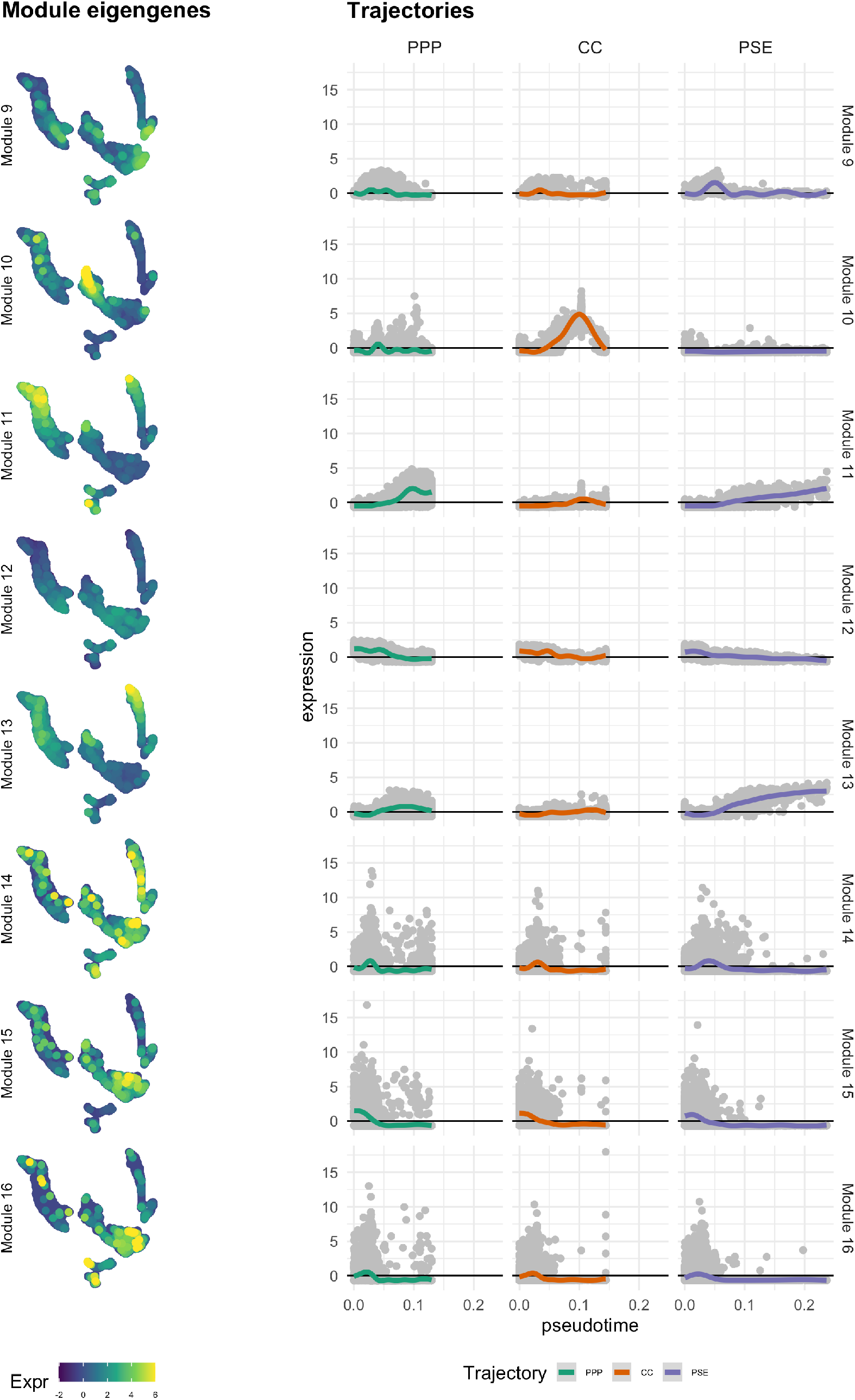
Gene co-expression network analysis identified 16 gene modules, represented by eigengene profiles and their expression along PPP, CC and PSE trajectories. Gene expression of each module is summarised by the eigengene profile, which is the first principal component score from a PCA done on the expression matrix and are plotted on the UMAP and different trajectories to visualise module expression across various cell types and along developmental trajectories in the phloem pole. Module 1 consists of 1367 genes (with variance explained score 21.4%) and shows an increasing expression in both PPP and CC trajectories, while lower than average expression in PSE. Modules 2, 3 and 7 contain 995, 878 and 225 genes respectively, which are highly expressed in the early cells. Module 4 with 778 genes shows broad expression in CC, PPP and PSE, but was lowly expressed on all three trajectories. Module 5, containing 368 genes, shows expression in cell clusters 12 and 13 while the 291 genes in Module 6 are mostly expressed specifically for PSE cells. The 164 genes in Module 8 were mostly expressed in the outer layers, with some lowly expressed in the mature PPP and CC cells. The eigengene profile of Module 9 (134 genes) shows expression in all cell types except CC, but particularly higher in early cells while Module 10 contains 18 genes highly expressed in CC. Modules 11-16 contain no more than 10 genes and the exact sizes are 6, 5, 3, 2, 2, and 2 genes respectively.

**Fig S7.**
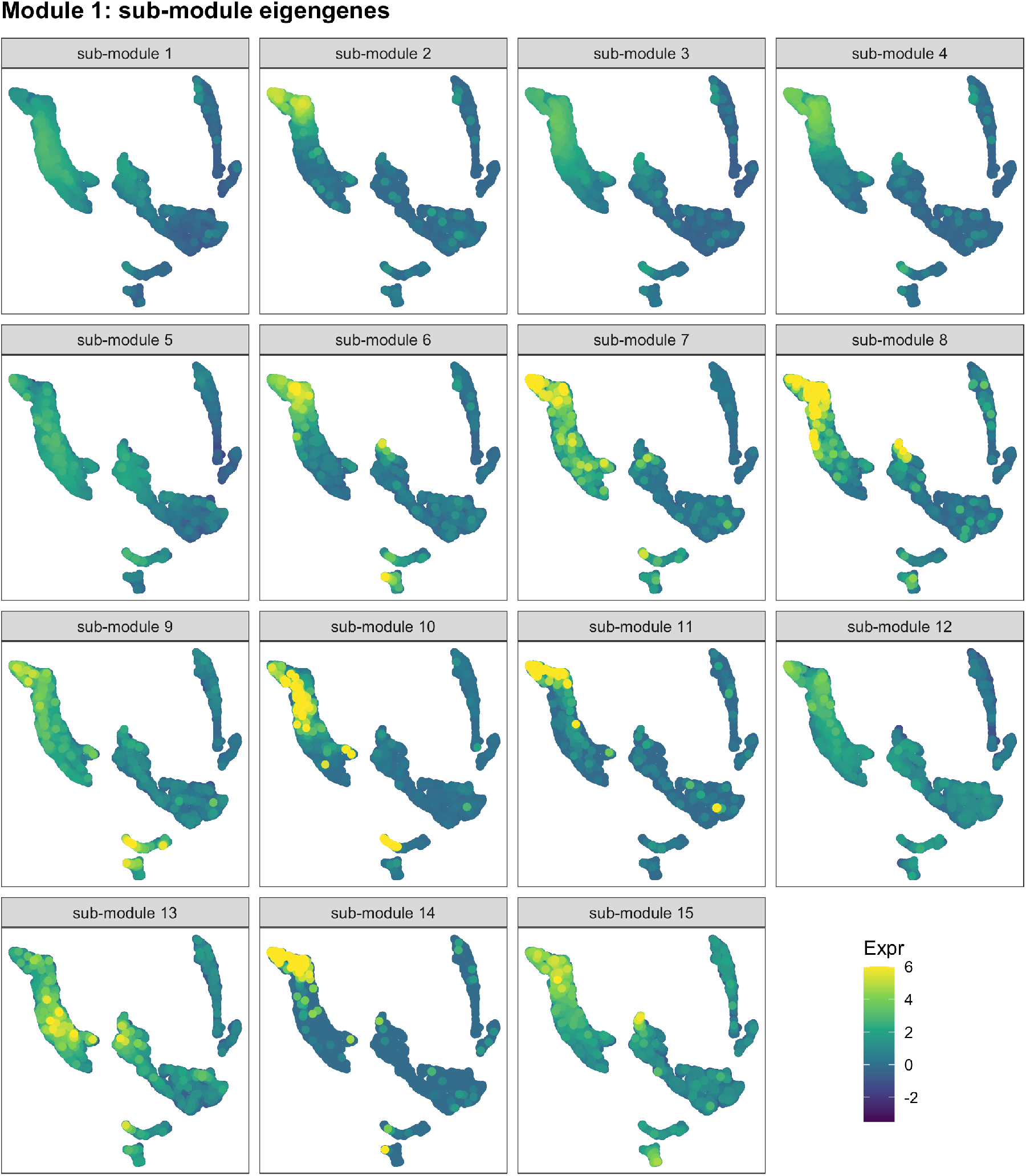
Analysis on sub-network of Module 1 genes identified 15 sub-modules via Louvain algorithm, showing expression patterns represented by sub-module eigengene profiles. The eigengene profile of sub-module 1, containing 326 genes, shows expression in both PPP and CC with relatively low expression in PSE, similar to the pattern revealed by Module 1 eigengene. 8 out of 9 genes with ring-specific expression pattern found in Module 1 fall in this sub-module, while *At3g16330* falls in sub-module 3, which shifts slightly towards mature pericycle cells. Sub-module 2 contains 318 genes specifically for pericycle cells. Genes in sub-module 4 and 6 are highly expressed in PPP cells and some in out layers. For sub-module 5 and 7, the eigengene profiles show relatively broader expression in both PPP and CC, as well as the out layers. The other 8 sub-modules contain no more than 9 genes.

**Fig S8.**
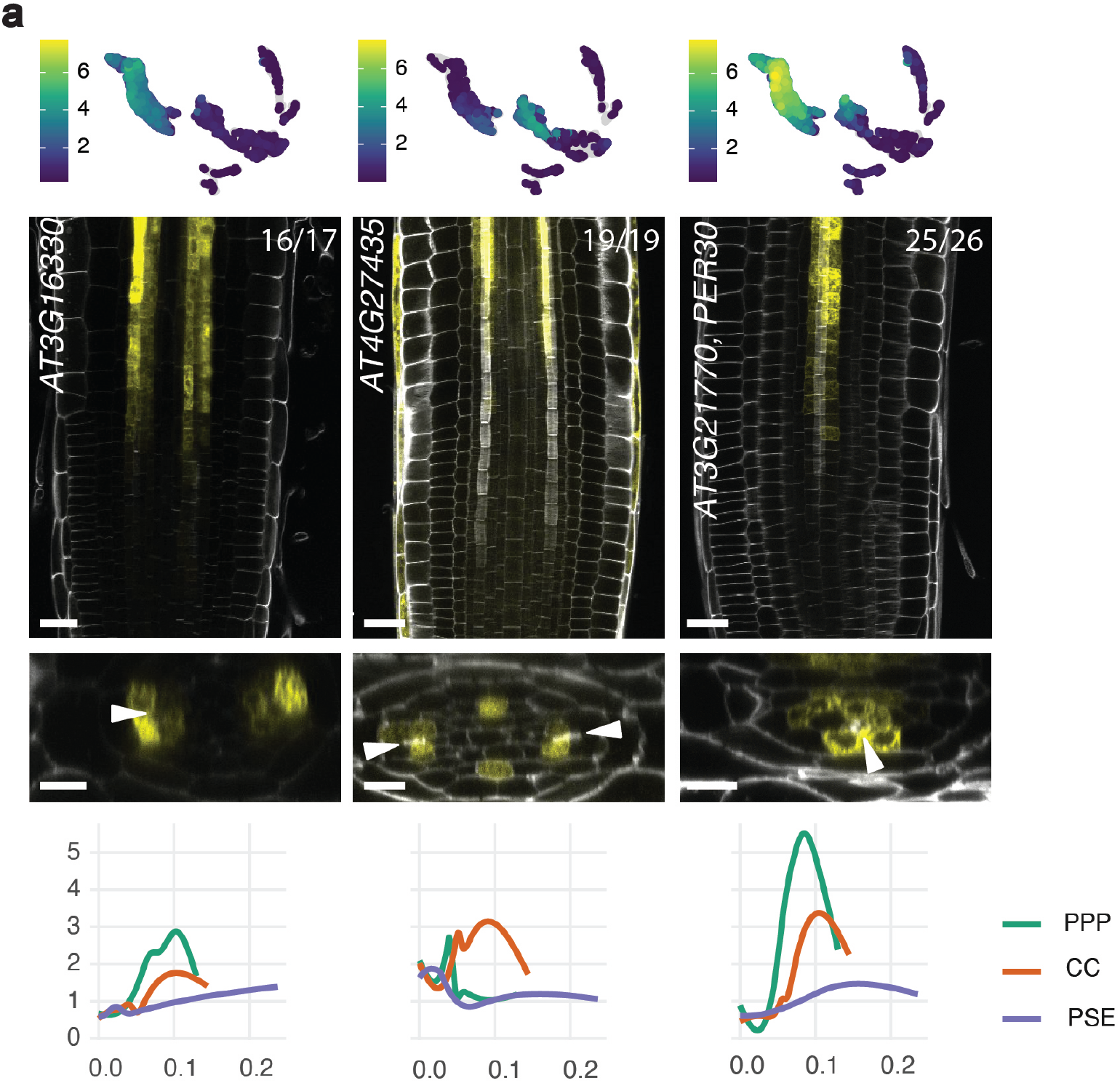
Module 1 also groups genes with an extended or partial ring expression pattern. a) New genes with an expression pattern validating the module eigengene analysis. All the genes presented in this panel are expressed forming a ring pattern (*At3g16330*(cite Miyashima) or extended ring (*At4g27435*, note the expression in protoxylem, and *PER30*, note the expression in procambium) at the time of PSE enucleation. They are all grouped in module 1, except *At4g27435*, which belongs to module 4. UMAPs show the particular cluster-weighted normalised expression of each gene in the phloem pole cell atlas and microscopy pictures are representative images of the transcriptional reporter lines where the gene promoter is fused to *VENUSer*. Scale bar in the longitudinal sections is 25 µm while it is 10 µm in the cross sections. White arrowheads point to PSE cells as a reference point. Each gene has also been plotted in PPP (green), CC (orange) and PSE (purple) trajectories, showing average expression values in the Y-axis and pseudotime in the X-axis. The numbers in each panel indicate samples with similar results, of the total independent biological samples observed.

**Fig S9.**
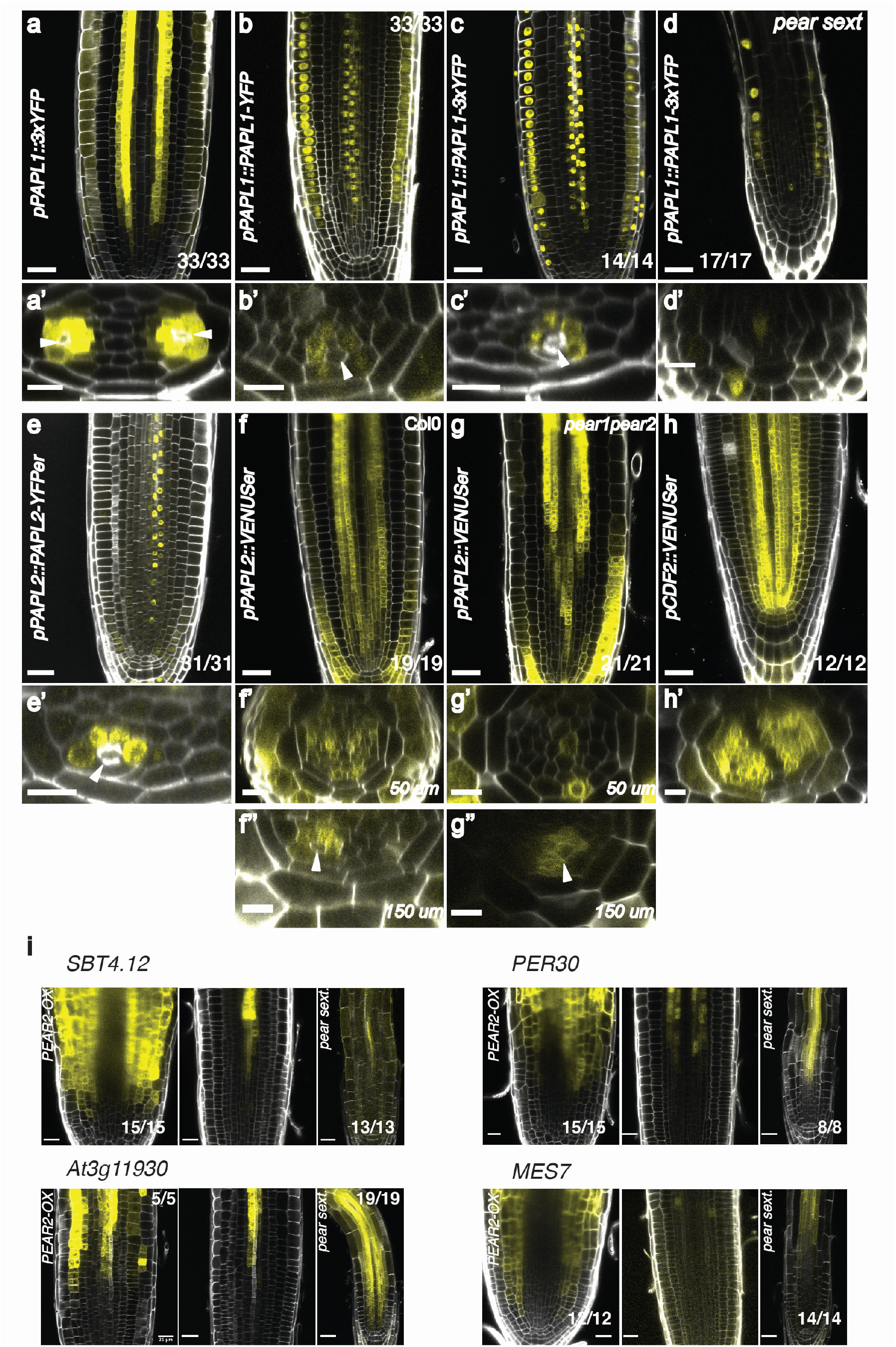
Detailed analysis of *PAPL* gene expression. a) *pPAPL1::3xYFP* showing a strong expression in all the cells surrounding PSE and a weaker expression in the neighbouring procambial layer. The latter is only observed with the 3xYFP reporter. b) Nuclear localisation of *pPAPL1::PAPL1-YFP* in epidermis and PSE adjacent cells which recapitulates the 3xYFP fusion pattern (c), indicating that PAPL1 is not mobile. Occasionally some nuclei appear highlighted in the neighbouring procambial layer, where this gene is expressed weakly as shown in a. d) Phloem meristematic expression of *pPAPL1::PAPL1-3xYFP* disappears in pear sext. while the epidermis signal stays. Occasionally the reporter was also observed in a central xylem cell. e) *pPAPL2::PAPL2-YFP* recapitulates *PAPL1* translational expression pattern. f) *pPAPL2::YFPer* expression mirrors the *PAPL1* ring transcriptional expression, although it has a broader domain close to QC and it is expressed in columella and epidermis. g) Like *PAPL1*, *PAPL2* ring expression domain gets delayed in *pear1pear2* mutant. h) Transcriptional reporter lines where the promoter was fused to *VENUSer* were transformed into *pear sext* mutant background and *pRPS5A::PEAR2-GR*, a line overexpressing ectopically *PEAR2* in the whole meristem. *PEAR2* was sufficient to induce the expression of the different genes in these layers. Primed letters show the cross section of each respective letter. Scale bar in the longitudinal sections is 25 µm while it is 10 µm in the cross sections. White arrowheads point to PSE cells as a reference point. µm in the cross sections indicate the distance from QC. The number in each confocal picture indicates samples with similar results of the total independent biological samples observed.

**Fig S10.**
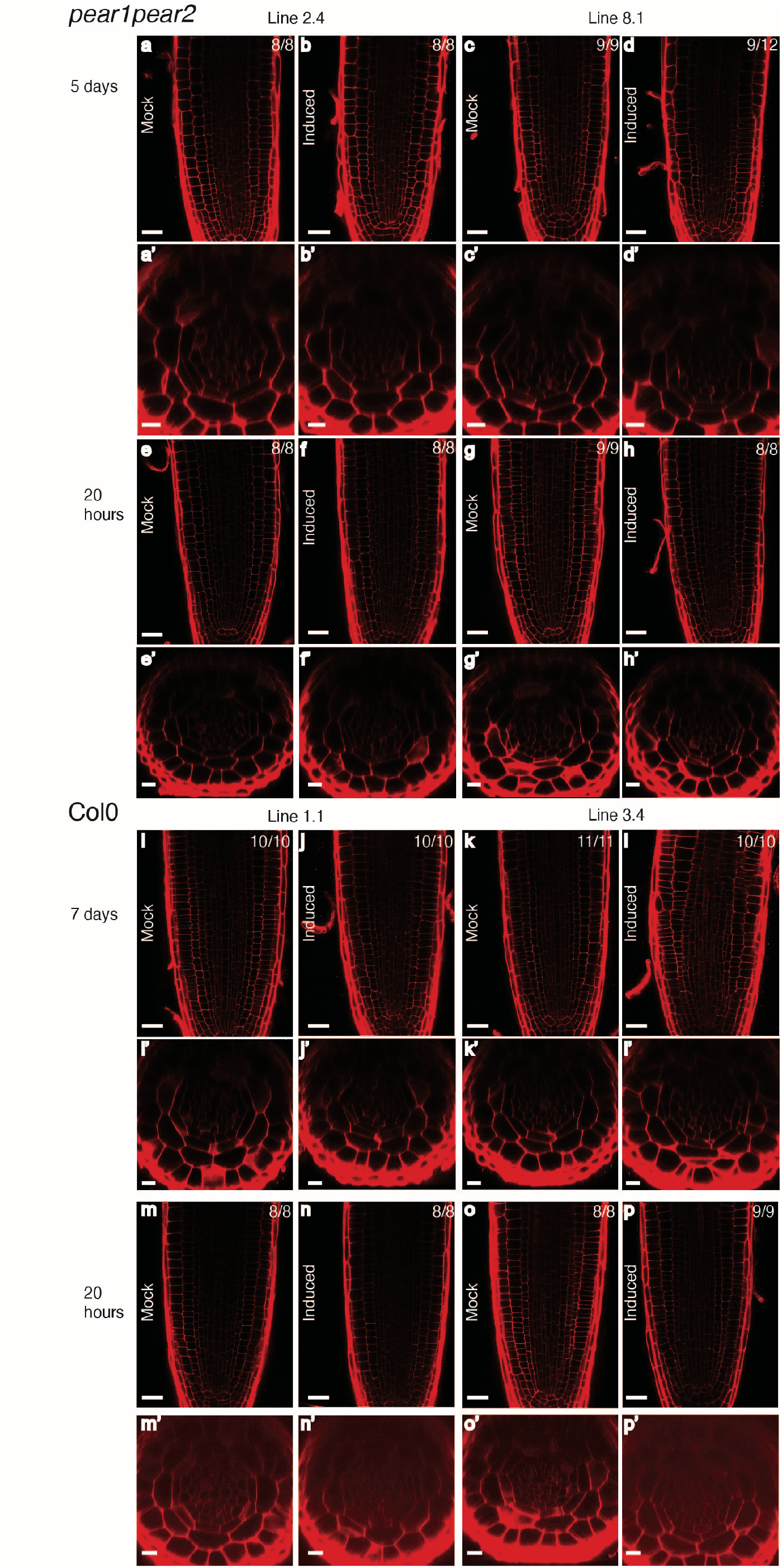
*PAPL* genes do not induce periclinal cell divisions. See next page for caption. 2 different lines for *pWOL::XVE>>PAPL1* in *pear1pear2* mutant background were induced in beta-estradiol for either 5 days (a-d) or 20 h (e-h). The same construct was transformed in the Col0 background, with 2 lines carried forward. Seedlings were induced for either 7 days (i-l) or 20 h (m-p). In the mock treatment, DMSO was added to the media instead of beta-estradiol. Primed letters show the cross sections of each respective letter. Scale bar in the longitudinal sections is 25 µm while it is 10 µm in the cross sections. The number in each panel indicates samples with similar results of the total independent biological samples analysed.

**Fig S11.**
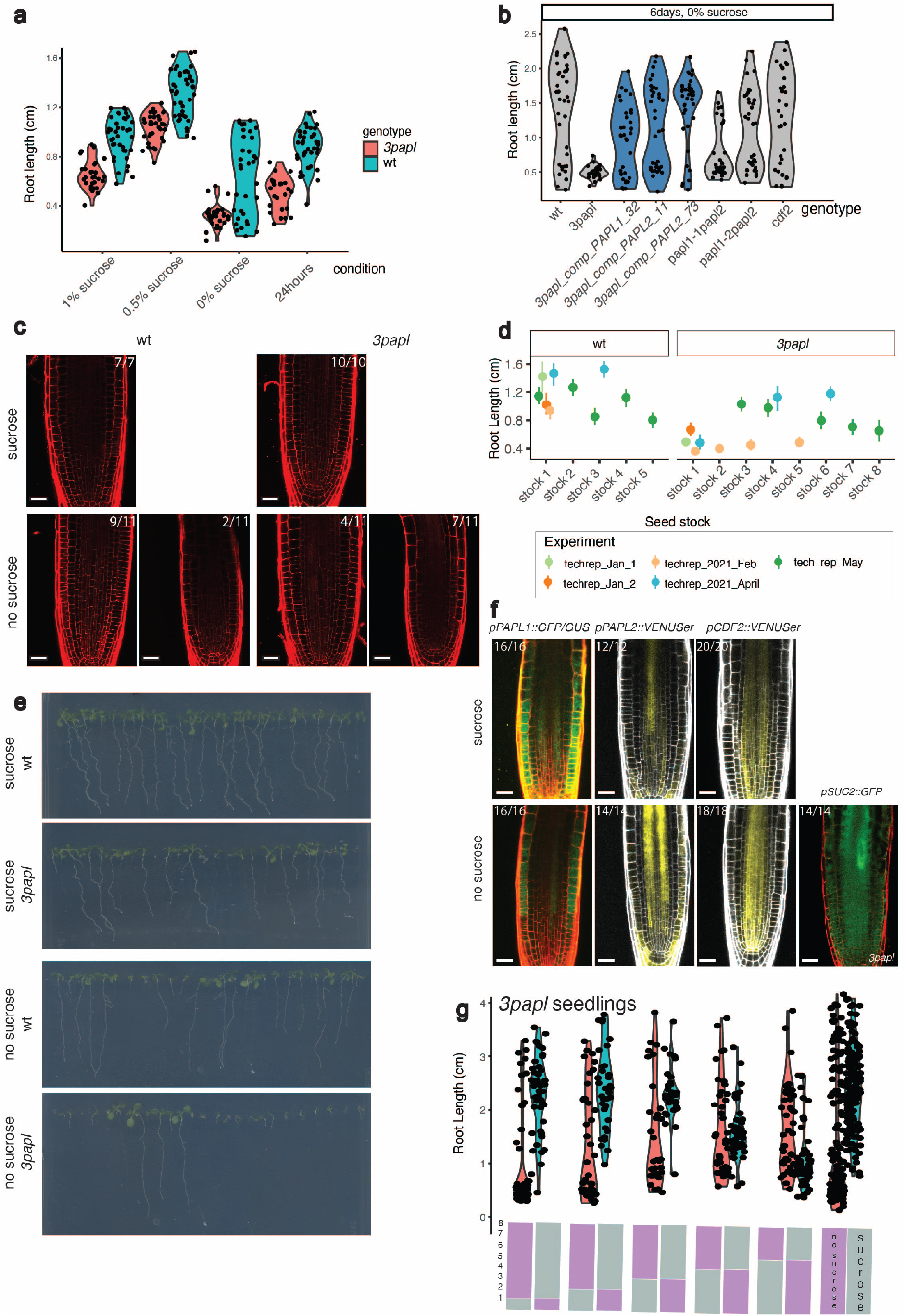
*PAPL* genes seem to be important for a correct root nutrition. See next page for caption. a) Root length in cm for 5dps *3papl* and wt seedlings in different conditions (1% sucrose, 0.5% sucrose, 0% sucrose and 24-hour light regime). For 0.5% sucrose, 42 wt and 36 *3papl* seedlings were used. For 1% sucrose, 39 wt and 29 *3papl* seedlings, for 24 hours, 41 wt and 25 *3papl* and for 0% sucrose, 34 wild type and 30 *3papl* seedlings b) Root length in cm for 6 dps seedlings of wt, *3papl* mutant, 3 complementation lines, double mutants and *cdf2* single mutant grown in media depleted of sucrose. 36 seedlings were measured for wt, 37 seedlings for *cdf4cog1-7*, 33 for *cdf4cog1-6*, 32 for *cdf2*, 28 for *3papl*, 38 for *3papl complementation PAPL2 line 7.3*, 37 for *3papl* complementation *PAPL2 line 1.1* and 34 for *3papl complementation PAPL1 line 3.2* c) Confocal pictures of 7 dps wt and *3papl* seedlings grown in sucrose containing media or media without sucrose d) Same data as in Fig 4g, but showing the variation across experiment and seed stock batches (only wt and 3*papl* are shown for illustration, but similar variation was observed for the complementation lines). Mean and 95% confidence interval per experiment were estimated by bootstrap (500 samples). e) Confocal pictures of 3 dps roots expressing reporters for *pPAPL1::GFP/GUS*, *pPAPL2::VENUSer*, *pCDF2::VENUSer* in Col0 background or *pSUC2::GFP* in *3papl* background in media containing/depleted of sucrose f) Scans of 8 dps seedlings (wt, stock 1, and *3papl,* stock 8) grown in 1% sucrose media or without sucrose. The number in each confocal picture indicates samples with similar results of the total independent biological samples analysed. g) Replicate of the transfer experiment between sucrose and sucrose-depleted plates of *3papl* seedlings (stock 8). Time (in days) spent in sucrose and without sucrose is represented by a grey and purple bar respectively. Transfer was done days 1-5 and all roots were measured at 8 dps. The bars are divided in 8 portions representing the days in each condition. 203 seedlings were grown as a control without sucrose, 156 were grown as a control with sucrose and a total of 99 seedlings were transferred on day 1, 75 on day 2, 70 on day 3, 93 in day 4 and 96 in day 5.

**Fig S12.**
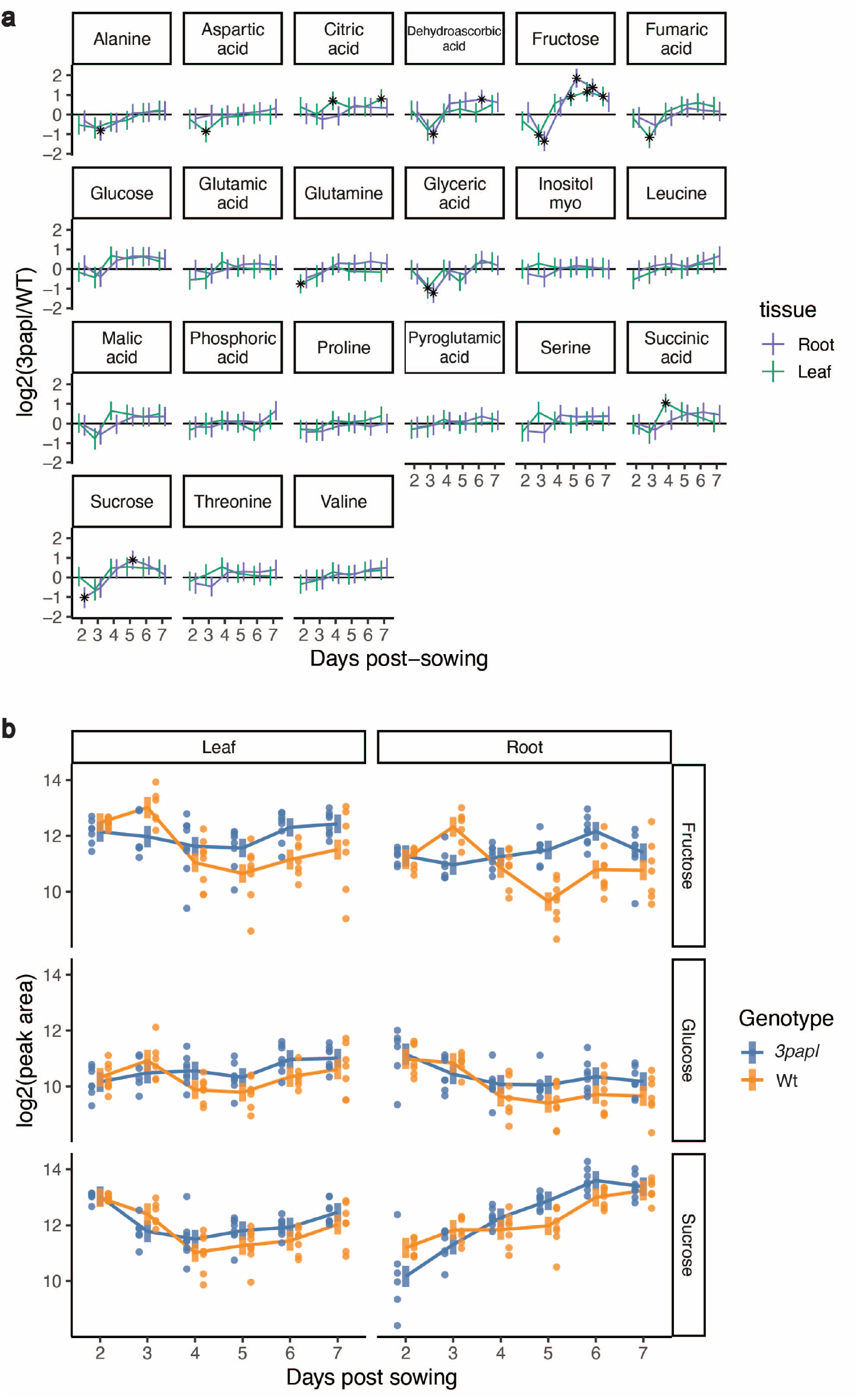
Difference in WT and 3papl metabolite levels in leaves and roots. a) Overview of the different metabolites with significant differences between WT and mutant in at least one of the time points. The error bars show 2 times the standard error (i.e. an approximate 95% confidence interval) estimated from our linear model (see methods). The asterisk highlights points that were statistically significant after adjusting for multiple testing across all the tests (false discovery rate of 5%). b) Average metabolite levels for sucrose in mutant and wild-type. The bars denote the 95% confidence interval estimated from our linear model (see methods). The points show the raw data for individual samples.

## Notes

https://www.ncbi.nlm.nih.gov/geo/query/acc.cgi?acc=GSE181999

https://www.ncbi.nlm.nih.gov/geo/query/acc.cgi?acc=GSE182672

https://doi.org/10.17863/CAM.74836

https://github.com/tavareshugo/publication_Otero2021_PhloemPoleAtlas

### Competing Interest Statement

The authors have declared no competing interest.

